# Phenotypic and molecular convergence of a cancer-associated pigmentation trait in *Xiphophorus evelynae*

**DOI:** 10.64898/2026.11.25.769034

**Authors:** Sophia K Haase Cox, Nadia B Haghani, John J Baczenas, Theresa R Gunn, Tristram O Dodge, Kelsie E Hunnicutt, Alexandra E Donny, Andreas Rosenwald, Alexa Pollock, Daniel L Powell, Paola Fascinetto-Zago, Manfred Schartl, Stepfanie M. Aguillón, Molly Schumer

## Abstract

Cases of phenotypic convergence are common across the tree of life, and often reflect shared ecological pressures. Studying such cases can provide insights into the repeatability of evolution in nature and help uncover the degree to which similar morphological phenotypes tend to also be driven by convergent molecular mechanisms. Here, we investigate the genetic basis of a convergent phenotype in *Xiphophorus*, a genus of fish known to possess a wide variety of polymorphic pigmentation traits, which have been shown to play important roles in their behavioral and ecological environments. We investigate a melanic spotting polymorphism occurring in the caudal fin that we call “confetti caudal” in *X. evelynae.* This trait is phenotypically similar to tailspot polymorphisms in two other *Xiphophorus* species, diverged from *X. evelynae* by ∼6 million generations. We characterize the developmental progression of this trait and use a genome-wide association study paired with analysis of long-read sequencing data to narrow in on a region containing a structural variant that differs between spotted and unspotted fish. Analysis of RNA sequencing data shows that *xmrk*, a known pigmentation gene that falls within this region, is differentially expressed. Notably, *xmrk* has been implicated in the development of tailspots in deeply diverged *Xiphophorus* species. We suggest that *xmrk* is responsible for confetti caudal in *X. evelynae* and has been repeatedly co-opted to drive tail patterns across the genus. Despite confetti caudal segregating at high frequency within *X. evelynae,* we detect a number of individuals in which the spotting pattern progresses into melanoma. Overall, our results highlight convergent involvement of the same gene in tailspot traits in two distantly related lineages. However, we also find that patterns of association between the tailspot pattern, *xmrk,* and melanoma vary between species, highlighting the complexity of convergent evolution at the molecular level, with implications for how convergent traits emerge and evolve.

## Introduction

A long-standing debate in the fields of evolutionary genetics and developmental biology is whether evolution is predictable at the genetic level—researchers have disagreed on whether “replaying the tape of life” would produce similar results at different biological scales (Gould 1989; Orgogozo 2015). Cases of recurrent or convergent evolution provide a powerful system to begin to answer some of these fundamental questions. At the morphological level, evidence of convergent evolution is abundant (Cerca 2023; Losos 2011; Stern 2013) and can provide insight into the ecological forces driving trait evolution. Repeated independent evolution of similar phenotypes may point to shared selective pressures driving their evolution and maintenance, so studying such cases of convergence can be particularly valuable when the function of the trait is unknown.

One open question is whether similar phenotypes that have evolved independently in distinct lineages tend to be convergent at the molecular level. Such molecular convergence may involve repeated co-option of the same genetic pathways (Bilandžija *et al*. 2012; Chan *et al*. 2010; Moreno *et al*. 2024), repeated changes in the same genes (Miller *et al*. 2007; Toprak *et al*. 2012), or even repeated changes to the same positions in the coding sequence (Dobler *et al*. 2012; Labeyrie 2003; Rosenblum *et al*. 2014; Stern 2013; Zhen *et al*. 2012). Until recently, this question was exceptionally challenging to address in most species. However, increasing availability of whole genome sequencing of diverse species has made addressing this question newly tractable, opening the door to compare the genetic mechanisms underlying convergent phenotypes. Empirical studies to date have been mixed, with some studies uncovering evidence for repeated co-option of the same genes or genetic pathways, as well as evidence from others of distinct genetic mechanisms (Cerca 2023; Christin *et al*. 2010; Conte *et al*. 2012; Rosenblum *et al*. 2014; Stern 2013; Tenaillon *et al*. 2012). Moreover, these outcomes may shift as a function of evolutionary history or the genetic architecture of the trait in question (Christin *et al*. 2010; Elkin *et al*. 2023; Milligan & Sella 2026; Peichel & Marques 2017).

Pigmentation traits are particularly tractable for studying molecular and phenotypic convergence because they are easy to observe, can have clear fitness consequences, and tend to have simpler genetic architectures (Kratochwil & Mallarino 2023; Kronforst *et al*. 2012; Orteu & Jiggins 2020). Black spots on the caudal (tail) fin of fish are one such pigmentation trait that has convergently evolved across deeply diverged species. These tail spots are found broadly across ray-finned fish lineages, varying in their exact presentation, but in most cases their function is unknown (Giammona 2021). Many species have a visually similar single black spot on the base of the caudal fin (McPhail 1977; Nursall 1973; Winemiller 1990), while other species have multiple spots speckled across their caudal fins (Kittilsen *et al*. 2009; Rodriguez-Silva *et al*. 2023; Thomas *et al*. 2007). Researchers have speculated that these patterns serve as deflection marks to reduce fatal injury from predators, or as a visual cue involved in intraspecific communication in schooling species (McPhail 1977; Nursall 1973; Winemiller 1990). The broad distribution of tailspots across deeply diverged lineages suggests many independent evolutionary origins of spots. As one example, eyespot bowfins (*Amia ocellicauda*) are about 320 million years diverged from teleost species with phenotypically similar spots (Dornburg *et al*. 2014; Hughes *et al*. 2018; Hurley *et al*. 2007). Black tailspots are found in fish that occur across dramatically different environments, predation pressures, and propensity for schooling, hinting that multiple mechanisms may drive the evolution of these traits in different species.

While tailspots have arisen recurrently across deeply diverged species of fish, there are also cases of close relatives where tailspots appear to have arisen independently. These cases present a more tractable path forward in assessing shared genetic architecture and evidence for shared functional consequences of tailspots. Swordtails and platyfish in the genus *Xiphophorus* are an excellent model for studying the genetic basis of pigmentation traits (including tailspots) due to high within and between species pattern variability (Culumber 2014). Melanic traits in *Xiphophorus*, like tailspots, are caused by one of two pigment cell types: micromelanophores or macromelanophores. Micromelanophores contribute to gray body color in *Xiphophorus* fish, and also several body and tailspot pigment patterns (Basolo 2006; Cui *et al*. 2013). Macromelanophore pigment cells are much larger than micromelanophores, and are visible as large dark blotches of pigment on the body or fins (Anders *et al*. 1979; Gordon 1948; Kallman & King 1975; Wellbrock *et al*. 2002). Several *Xiphophorus* species are polymorphic for macromelanophore tailspot patterns collectively referred to as “spotted caudal,” including *X. cortezi* and *X. birchmanni* (Culumber 2014; Kallman 1972). While the impacts of many of these patterns are unknown, melanin pigmentation patterns can play both an ecological and social role in *Xiphophorus* (Franck et al. 2001). Some studies have found relationships between melanism and behavioral differences such as boldness (Culumber 2016), as well as associations with ecological traits such as thermal tolerance (Culumber *et al*. 2014; Payne *et al*. 2022). *X. cortezi* males with spotted caudal have been shown to be more aggressive to conspecifics (Fernandez 2010) and preferred by females (Fernandez & Morris 2008), suggesting that this trait may confer a competitive or sexually-selected advantage. Despite these potential advantages, macromelanophore patterns in *Xiphophorus* are also associated with the development of malignant melanoma (Schartl *et al*. 1995), particularly in interspecific hybrids. It has been known since the 1930s that hybrids between *X. maculatus* and *X. hellerii* develop melanomas emerging from macromelanophore pigmentation spots (Lu *et al*. 2020; M Gordon 1937; Walter & Kazianis 2001), driven by the *xmrk* (*Xiphophorus* melanoma receptor kinase) gene. More recently this result has been replicated for hybrids between other species (Powell *et al*. 2020).

Here, we investigate the genetic architecture of a macromelanophore tailspot polymorphism in the platyfish *X. evelynae*. We refer to this trait, whose genetic basis has not been previously studied, as “confetti caudal”. This trait is visually similar to the macromelanophore tailspot found in *X. birchmanni* and *X. cortezi* (“spotted caudal”), but occurs in *X. evelynae*, which is ∼6 million generations diverged from the clade containing *X. birchmanni* and *X. cortezi* (Du *et al*. 2024). We used large population samples paired with a genome-wide association study to investigate the genetic basis of confetti caudal. Despite the substantial evolutionary divergence between *X. evelynae* and the lineages leading to *X. cortezi* and *X. birchmanni*, we find the confetti caudal phenotype is associated with *xmrk.* We investigated the developmental timing of the trait in juveniles and adults and found evidence of spontaneous melanoma formation in *X. evelynae*. Our results provide new insights into how the same gene can be recurrently co-opted to generate a convergent pigmentation phenotype.

## Methods

### Sample collection and phenotyping

Sampling from the wild-caught population of *X. evelynae* was previously collected for the study of a different pigmentation trait (see Haghani et al. 2025). We collected *X. evelynae* individuals from the Juntas Chicas population in 2023 using baited minnow traps. This site is on a tributary of the Río Tuxpan directly upstream of the town of Juntas Chicas in the state of Hidalgo, Mexico (20.573257° N, 98.031427° W). Fish were anesthetized in a buffered solution of 100 mg/mL MS-222 and river water, and standardized photographs were taken on both sides of the body with caudal and dorsal fins spread. A small tissue sample was taken from the caudal fin of each individual for genome sequencing. Fish were allowed to recover in river water before being released back into the population. In total, 262 fish were sampled, including males (n = 56), females (n = 130), and juveniles (n = 76). Permission to sample fish was provided by the Mexican government (Permit No. PPF/DGOPA-002/19) and followed procedures described in Stanford Laboratory Animal Care protocol #33071.

We used these standardized photos for individual phenotyping and explored a variety of phenotyping approaches (**Supplementary materials 1, 2**). A categorical approach based on the intensity and number of spots (**Fig. S1)** was used for most analyses, while a quantitative method (**Supplementary materials 1, Fig. S2**) was used for tracking the progression of the confetti phenotype in adults. We found strong correlation between measurements using these two approaches (**Fig. S3**, Spearman rank correlation = 0.94, p < 0.0001). We evaluated whether confetti caudal occurs at different frequencies based on sex using a Pearson’s Chi-squared test with Yates’ continuity correction in R version 4.1.0.

### Measurement of confetti macromelanophores

In order to investigate if the confetti trait is made up of macromelanophores, we measured the size of the melanin cells in adult *X. evelynae* individuals with confetti caudal. Four *X. evelynae* females with confetti caudal were euthanized in an ice bath (following Stanford APLAC protocol #33071). The caudal fin was removed and mounted on a glass microscope slide (Fisherbrand ColorFrost Plus) in PBS (1x, 7.4pH) solution, immobilized with a cover slip, and imaged on a BioTek Cytation 5 imaging reader in color brightfield channels. Melanophores were measured using a protocol adapted from the procedure described in Given *et al*. (2026). Briefly, three melanophore cells from within the confetti pattern and three melanophore cells spatially separated from the confetti spot from each individual were measured using FIJI (Schindelin *et al*. 2012). Three measurements per cell were averaged to estimate the diameter of each cell due to the irregular shape of melanophores. Because macromelanophore cells are often overlapping, we only measured isolated macromelanophore cells on the edge of the pigmentation spots. This may bias us towards measuring smaller macromelanophores, but we expect this approach will be conservative. We evaluated whether there was a significant difference between melanophore cell size within versus outside the confetti spot with a linear mixed-effects model using the R package lme4 (Bates *et al*. 2015), including individual as a random effect variable.

### Tracking development of confetti caudal in juveniles

In order to investigate the developmental timing of the confetti caudal phenotype, we tracked lab-born *X. evelynae* fry over time by reanalyzing previously collected data (Haghani *et al*. 2025). Fry were collected within a day of birth from a tank of wild-caught *X. evelynae* adults from Juntas Chicas and immediately separated and housed individually in 2 L tanks with gravel substrate. We tracked a total of 32 individuals, with 30 surviving the full duration of the experiment. Photos were taken weekly for a period of 6 months. Each week, fish were anesthetized in MS-222, photographed on both sides, and examined under a dissecting microscope. We recorded the presence of confetti caudal spots as follows: fish with no visible macromelanophore spots on the tail were scored as 0, fish with some macromelanophores visible under the dissecting microscope were scored as 0.5, and fish with at least 1 melanin spot large enough to be visible without magnification were scored as 1. We additionally recorded the presence of other melanin pigmentation patterns and the stage of gonopodium development in males. After approximately 5 months, fish were individually labeled by tagging with a unique elastomer tag combination, housed together, and a final photograph was taken after one additional month. We also compared the developmental timing of confetti caudal in *X. evelynae* to that of the spotted caudal tail spot in *X. birchmanni* (**Supplementary materials 3**).

### Tracking changes in confetti caudal presentation in adults

Among adults with confetti caudal we observed a wide variation in the size and number of spots. This variation may be due to genetic or environmental factors, or it may simply reflect the age of the fish if the pattern intensifies over time. To determine if confetti caudal spots continue to expand in size or number in adult fish, we tracked 12 adult lab-born *X. evelynae* with confetti caudal, descended from wild-caught fish. These adults were anesthetized using 100 mg/mL MS-222, photographed, given an individually identifiable elastomer tag, and re-photographed after 64 and 171 days. One fish in the tank we originally sampled from did not have confetti caudal at the beginning of tracking, but developed it during the course of the experiment, and thus we added this fish to our sampling for subsequent timepoints (tracking n = 13 fish). We used ImageJ and the R package recolorize (Weller *et al*. 2024) to determine the amount of the caudal fin covered in melanic spots (**Supplementary materials 1, Fig. S2**). We used a paired Wilcoxon signed-rank test to test for differences in the fraction of melanophore coverage between pairs of time points, excluding missing data.

### DNA extraction and Tn5 library preparation

Sampling of DNA from fin-clips of the wild-caught population of *X. evelynae* was previously collected for the study of a different pigmentation trait (see Haghani *et al*. 2025). Samples were extracted and prepared for whole-genome sequencing following the approach described in Payne et al. (2022). Briefly, DNA was extracted using the Agencourt DNAdvance magnetic bead-based purification system (Catalog #A48705, Beckman Coulter, Brea, California). The manufacturer’s protocol for DNA extraction from tissue was followed except that half reactions were used. Tagmentation-based libraries were prepared for low-coverage whole genome sequencing using the Illumina DNA TDE1 Enzyme and Buffer Kit (Illumina, San Diego, CA). The libraries were bead purified using homemade 18% SPRI beads, quantified using a Qubit 4 Fluorometer (Thermofisher Scientific, Wilmington, DE), and the library size distribution was visualized with a Tapestation 4200 (Agilent Technologies, Santa Clara, CA). Low-coverage (∼1x) whole genome sequence data were generated for these libraries using an Illumina HiSeq 4000 at Admera Health Services (South Plainfield, NJ).

### Genome wide association study (GWAS)

To identify genomic regions associated with the confetti caudal trait, we performed a GWAS using the phenotypic scoring described above and whole-genome sequencing data. We grouped individuals into cases (confetti scores 3-5, n = 81) and controls (confetti score 0, n = 90) based on their categorical score for the confetti spotting pattern (**Fig. S1**). Out of concern that ambiguous phenotypes may add noise to the data, we only retained individuals that clearly had the trait and those that clearly lacked it in this analysis, excluding individuals with confetti score 1 or 2 (resulting in n = 171 individuals). Reads were aligned to the *X. evelynae* reference genome (Haghani *et al*. 2025) using bwa (Li 2013). Mapped reads were sorted, quality filtered, and realigned around indels using samtools version 1.16.1 (Danecek *et al*. 2021; Li *et al*. 2009). Allele frequency differences between the case and control groups were estimated using the legacy version of samtools. We removed variants that had fewer than 80X coverage in the dataset, or with a minor allele frequency below 0.05.

To identify the appropriate genome-wide significance threshold, we shuffled individual phenotype labels and re-ran the full GWAS (171 individuals), repeating this procedure 500 times. We filtered output using the same criteria we had applied to the real data. For each simulation, we recorded the minimum p-value, resulting in a total of 500 minimum p-values. To select our p-value threshold, we took the 5% quantile of this distribution, roughly corresponding to the p-value at which we expect a 5% false positive rate. In our analysis of the empirical data, only one region crossed this significance threshold, and we explored candidate genes in the region using tools in the tidyverse package (Wickham *et al*. 2019). We also evaluated whether there was differentiation between cases and controls genome-wide, i.e., evidence of population structure associated with phenotype (**Supplementary materials 4**).

In order to assess if the sex-determining region overlapped with the association for confetti caudal, we performed two additional GWASes. First, we performed a GWAS for sex using males and females as cases and controls, excluding juveniles (n = 123). Next, we repeated the case-control GWAS for confetti caudal with females only (n = 76), using the same reference genome for read alignment, but with the Y chromosome removed.

### Investigation of linkage disequilibrium and structural variation on chromosome 21

Due to the large width of the GWAS peak and its position near the sex determining region, we wanted to quantify linkage disequilibrium in the region. However, we lacked population-level data at high coverage with which to perform this analysis. As an alternative method, we used the low-coverage data to evaluate general patterns in this region. We first converted our low-coverage data into pseudohaploid calls by randomly sampling one mapped read at a given site per individual (as in Dodge *et al*. 2024). We next converted the “pseudohaploid” calls to PLINK format and used the ld command in PLINK v1.90 to calculate pairwise R2 for sites that fell within this region (Chang *et al*. 2015) and plotted average LD in windows over physical distance across the GWAS peak.

We next looked for evidence of structural variation on chromosome 21. We used minimap2 (Li 2018) to align chromosome 21, the putative X chromosome, from previously generated long read reference genomes of an *X. evelynae* individual with confetti caudal and one without this trait (Haghani *et al*. 2025). We used minimap with the settings recommended by the SVbyEye R package (Porubsky *et al*. 2025) and then visualized the alignment with this package. We used the Seqtk gap tool (Li 2026) to ensure neither reference genome had a gap in the region of interest.

Based on the results of the structural variant analysis, we further investigated coverage in this region using two confetti caudal and two wild-type individuals for which PacBio data was previously collected for another project (Haghani *et al*. 2025). We mapped these reads to the confetti caudal reference genome using pbmm2 (https://github.com/PacificBiosciences/pbmm2) with the CCS preset, filtered reads with a mapping quality less than 60, and visualized coverage in the structurally variable region in IGV (Robinson *et al*. 2011; **Supplementary materials 5**).

### Differential expression analyses using RNA sequencing

We identified a genome-wide significant GWAS signal that overlapped with a structural variant. Because this region contained 133 genes and the structural complexity in the region prevented finer scale mapping with GWAS, we utilized RNA sequencing to further evaluate genes that may be linked to the confetti trait. We selected eight wild-caught adult male *X. evelynae* from Juntas Chicas—four with confetti caudal spots and four without. All fish had been housed in the same tank in the lab for several months. Fish were photographed, then euthanized (following Stanford APLAC protocol #33071) and immediately placed in a tube of chilled RNAlater (Thermofisher Scientific). Samples were refrigerated in RNAlater for several days before dissection and extraction. From the fish with confetti caudal spots, a portion of caudal fin covered with macromelanophores was dissected out. These tissue samples were primarily from melanized areas, but due to the small size of the spots on some individuals, two of the dissections included some non-melanized tissue. From the same individual, a corresponding non-melanized fin portion was also sampled. Caudal fin tissue samples were collected from the same fin region of four unspotted fish for comparison. We extracted RNA using the Qiagen RNeasy Mini Kit following the manufacturer’s protocol, with the exception that Monarch pestles (New England Biolabs) were used to homogenize the tissue. To control for batch effects in downstream analysis, samples from each of the three experimental groups were processed as triplicates. RNA library preparation and Illumina sequencing were performed by Admera Health Services (South Plainfield, NJ) using the NEBNext Ultra II Directional RNA Library Prep Kit for Illumina (New England Biolabs).

After confirming that all samples had more than 20 million 150 bp paired-end reads, adapters were trimmed with the programs cutadapt (Martin 2011) and TrimGalore (https://github.com/FelixKrueger/TrimGalore), and sequence quality was assessed with FastQC (Babraham bioinformatics n.d.). Reads were pseudoaligned to the *X. evelynae* reference transcriptome (Haghani *et al*. 2025) with *kallisto* (Bray et al. 2016). We generated raw transcript counts with *kallisto* and then imported these into R to conduct differential gene expression analysis with the package DESeq2 (Love *et al*. 2014).

To analyze genes differentially expressed in melanized confetti caudal tissue, we performed three separate comparisons: pigmented tissue from a confetti fish vs. unpigmented tissue from a wild type fish, pigmented tissue from a confetti fish vs. unpigmented caudal tissue from the same fish, and unpigmented tissue from a confetti fish vs. unpigmented tissue from a wild type fish. For comparisons between wildtype and confetti fish, we used an experimental design that included batch (day of RNA extraction) and tissue type. For the comparison between pigmented and unpigmented areas on confetti fins, we used an experimental design that included individual and tissue type (batch was redundant with individual in this analysis). We used a local dispersion model and evaluated overall patterns of gene expression across samples using a PCA implemented in the R package PCAtools (Blighe & Lun 2023). PCA plots for tissue type were visually assessed for outliers; none were identified. We next ran the DEseq command to normalize gene counts by library size, estimate group dispersion, fit a negative binomial generalized linear model, and test for significant differences in expression between groups. We calculated Shrunken log-fold changes using the R package ashr (Stephens *et al*. 2023). We imposed a filter to remove genes that had fewer than ten reads in four or more samples (n = 6,485), leaving 18,266 genes.

We compared gene expression between fish with confetti caudal vs. fish without confetti caudal and also evaluated expression patterns in melanized tissue vs. non-melanized tissue. Genes were considered to be differentially expressed between groups at an adjusted p-value < 0.05. Expression results were visualized using the R package EnhancedVolcano (Blighe *et al*. 2024). We also performed weighted gene co-expression network analysis to identify clusters of co-expressed genes associated with the confetti phenotype (**Supplementary materials 6**).

### Investigation of possible melanoma in *X. evelynae* and hybrids

Several *X. evelynae* confetti caudal adults raised in the laboratory were observed to have an expansion of macromelanophore cells into the musculature of the upper or lower caudal peduncle. Since this is a phenotype typically associated with the development of melanoma, we used a histology approach to evaluate if these expanded spots showed evidence of progression to malignancy. Two *X. evelynae* males with apparent macromelanophore expansion into the caudal peduncle were euthanized in an ice bath (following Stanford APLAC protocol #33071). The caudal peduncle and caudal fin were fixed in 4% formalin solution for at least 72 hours. The samples were then washed in ethanol, embedded in paraffin, and serially sectioned at 5 µm thickness. The sections were stained with hematoxylin and eosin. Slides were scanned and visualized with the QuPath 0.7.0 software. Images were visually assessed for the progression stages of melanosis.

Past work has repeatedly linked the presence of macromelanophore traits to the development of melanomas in interspecific hybrids (Lu *et al*. 2020; Powell *et al*. 2020; Walter & Kazianis 2001). *X. evelynae* in the Juntas Chicas population are sympatric with only one other *Xiphophorus* species, a lineage of *X. malinche*. We generated F1 hybrids to test if hybridization between *X. evelynae* and *X. malinche* was possible, and if F1 hybrids showed evidence of changes in the macromelanophore patterns that may suggest misregulation or the potential for melanoma. We successfully produced artificial crosses between male *X. evelynae* with confetti and female *X. malinche* (**Supplementary materials 7**).

## Results

### Confetti caudal is polymorphic in the Juntas Chicas population

We found that confetti caudal is generated by dense clusters of melanic cells on the caudal fin **(Fig. 1B)**. Based on the cell sizes, these are predicted to be macromelanophores **(Fig. 1C, S4, Table S1)**. We analyzed a dataset of 262 wild-caught male, female, and juvenile *X. evelynae* fish to estimate frequency in the Juntas Chicas population. We found that the confetti caudal trait was present in about 66% of individuals (phenotype score ≥1, 65.6 ± 2.93%). The frequency of confetti caudal significantly differed between sexes (**Fig. 1D,E**; 74.6±3.81% of adult females and 50.0±6.68% of adult males; *χ*-squared = 9.67, df = 1, p = 0.00187).

**Figure 1:**
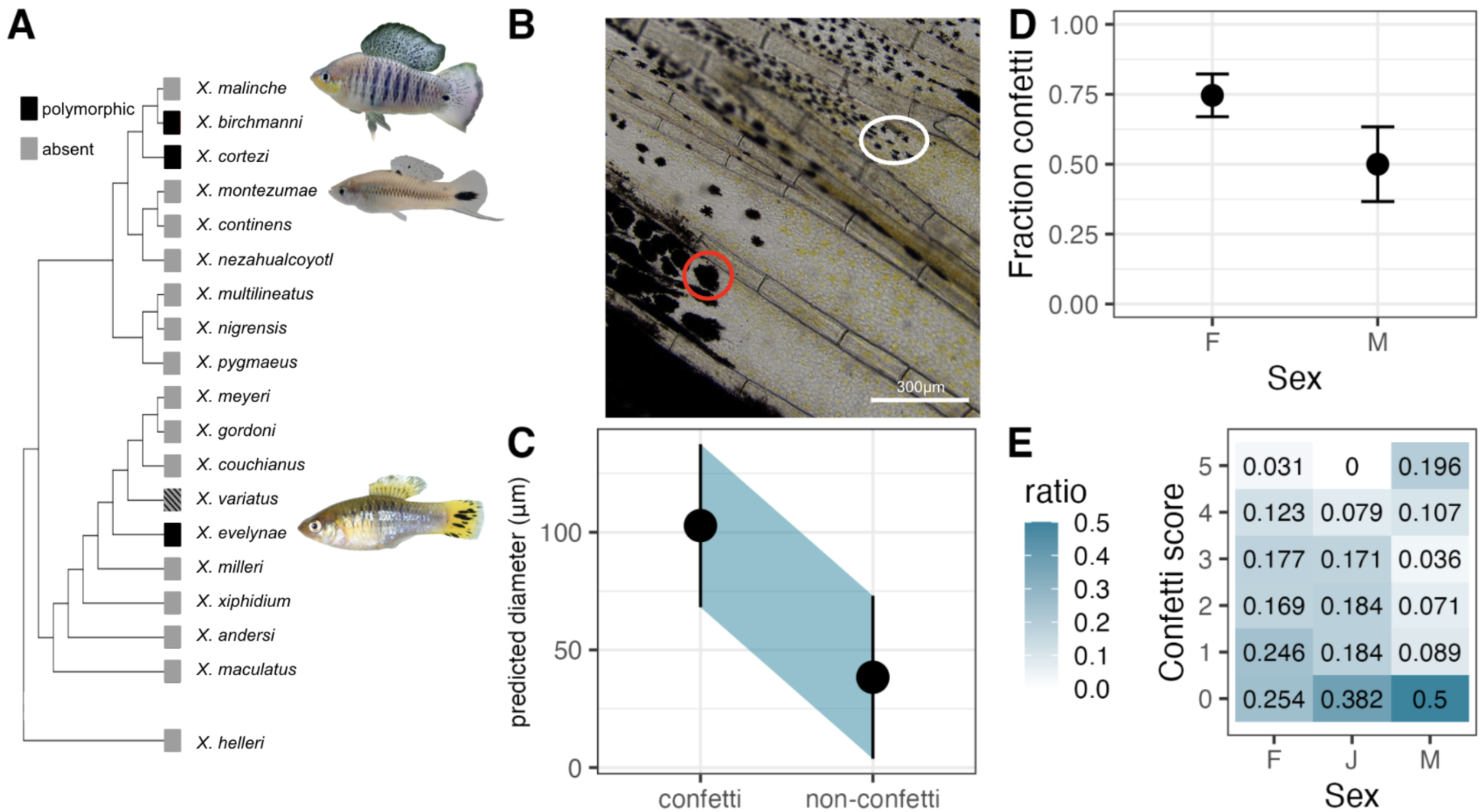
Confetti caudal is a polymorphic melanin tailspot trait in *X. evelynae*. **(A)** *Xiphophorus* phylogeny showing distribution of tailspot patterning. *X. birchmanni* and *X. cortezi* individuals with spotted caudal and an *X. evelynae* individual with confetti caudal are shown. It is unclear from the literature if *X. variatus* also possesses a macromelanophore tailspot pattern (**Supplementary materials 8**). **(B)** Image of the caudal fin of an *X. evelynae* individual with confetti caudal, showing the two types of melanophores found in *X. evelynae.* Red oval: macromelanophores as part of the confetti spotting pattern. White oval: micromelanophore cells not associated with the confetti phenotype. **(C)** Results of a linear mixed-effects model comparing the size of melanophores on the caudal fin of *X. evelynae* within the confetti trait vs. melanophores not associated with confetti caudal. Plotted is the predicted size of each melanophore category with the high and low confidence interval shown in blue (t-value =-4.56). **(D)** Frequency of confetti presence (score 1-5) in wild-caught male (n = 56) and female (n = 130) *X. evelynae* from Juntas Chicas (± 2 SE). Females have confetti at a significantly higher frequency than males (*χ*-squared = 9.67, df = 1, p = 0.00187). **(E)** Proportion of adult male (M), adult female (F), and juvenile (J) wild-caught *X. evelynae* with confetti scores 0-5 collected from the Juntas Chicas population.

### Timing of confetti caudal development in juveniles and trait progression in adults

To determine the age at which confetti arises in juveniles, we tracked the development of the trait in lab-born *X. evelynae* by following the onset of spot formation in individual fry (**Fig. 2A**). The age at which individuals first developed macromelanophores on the caudal fin was variable, ranging from 41 to 135 days post-birth. On average, juveniles developed macromelanophores on the caudal fin by 80.6 days, with a median age of development at 73 days old (n = 26). We also compared the timing of the onset of spot formation between *X. evelynae* and *X. birchmanni* males (**Fig. S5**). We found suggestive evidence that *X. evelynae* develops visible spots later than *X. birchmanni,* though not significantly so, potentially due to low sample size (**χ**-squared = 1.8591, df = 1, p = 0.1727) (**Supplementary materials 3**).

**Figure 2:**
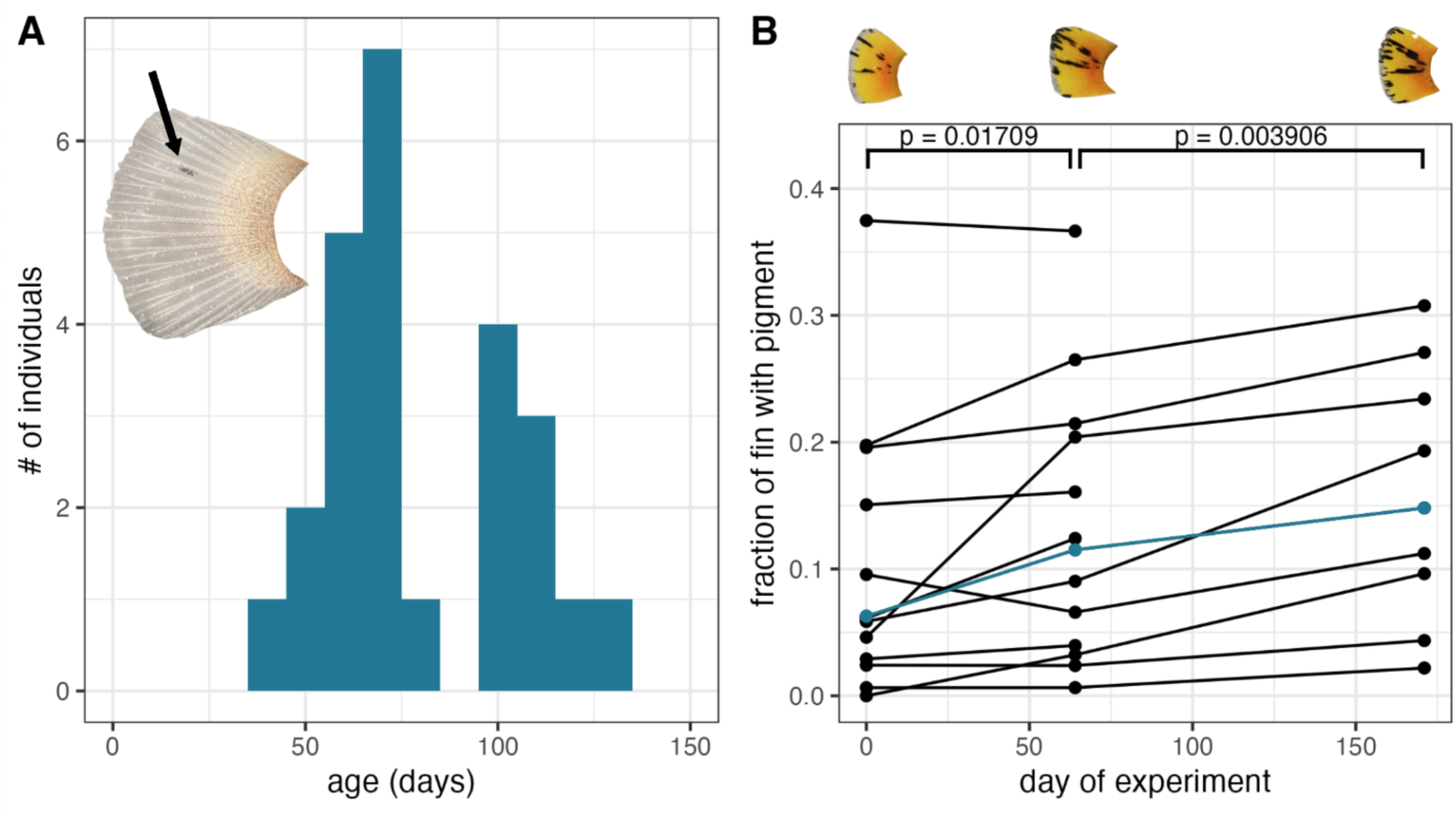
Development of confetti caudal in juveniles and adults. **(A)** Distribution of age (in days) at which juveniles first developed macromelanophores on the caudal fin based on visualization under a dissecting scope. Inset: Confetti caudal originates as one small macromelanophore spot. This representative photo was taken from a juvenile (age 109 days) at the first time point when confetti was detected under a dissecting scope. **(B)** Change in the fraction of the caudal fin covered by confetti spots in adult fish over time. Each line represents a different individual. Day 0 represents the first day photos were taken, and subsequent timepoints represent the number of days since that photo. The individual highlighted in blue corresponds to the caudal fin photos shown above the plot. There is a statistically significant difference in confetti score between the first and second time points (paired Wilcoxon signed-rank test: V = 7, p = 0.0171) and the second and third time points (V = 45, p = 0.00391).

To determine if the confetti caudal trait changes in presentation over time once it has developed, we tracked the proportion of the caudal fin that was pigmented in adult *X. evelynae* (all at least 1 year old at the start of the experiment). We found that even in adult fish, there is a high degree of variation in the number and size of macromelanophore spots, and that pigmentation on the fin increased over time (**Fig. 2B**). Specifically, we found a significant difference in confetti score between our first and second sampling time points (paired Wilcoxon signed-rank test: V = 12, p = 0.0171) and our second and third sampling time points (V = 45, p = 0.00391). Note that in one case measurements slightly decreased between the first and second timepoints, which we attribute to small differences in the spread of the fin from our photographs.

### A structural variant containing *xmrk* is associated with confetti caudal in *X. evelynae*

We used a genome-wide association study (GWAS) to identify regions of the genome associated with the confetti caudal phenotype. We identified a strong signal on the putative X chromosome, chromosome 21, which exceeded the genome-wide significance threshold determined via permutation (**Fig. 3A**). The significant interval is ∼4 Mb and overlaps with 133 genes, including the gene *xmrk* (**Fig. 3B**). The distal end of chromosome 21 is the sex chromosome in several *Xiphophorus* species, and we found the GWAS region overlapped with the sex determining region in *X. evelynae* (**Fig. S6**). Because including the Y chromosome variants in the GWAS might affect the results, we repeated the analysis using females only, and found a similar pattern despite reduced power (**Fig. S7**). We also confirmed there was no evidence that the confetti phenotype was associated with population structure in *X. evelynae* (**Fig. S8**). An alignment between the X chromosome of an *X. evelynae* with confetti caudal and the X chromosome of a wildtype fish shows evidence of an approximately 1 Mb structural variant in the center of the associated region (**Fig. 3C**). We confirmed that this presence/absence variant was associated with the presence or absence of the confetti phenotype in other samples for which we had access to long-read data (**Fig. S9A**). Consistent with this inference from long-read data, we find substantially reduced coverage based on short-read data in this region in unspotted versus spotted samples from our GWAS (**Fig. S9B**). Given that the SNPs most strongly associated with the confetti phenotype directly flank the structural variant, we were particularly interested in investigating the functions of the six genes found in this region (**Fig. 3C**).

**Figure 3:**
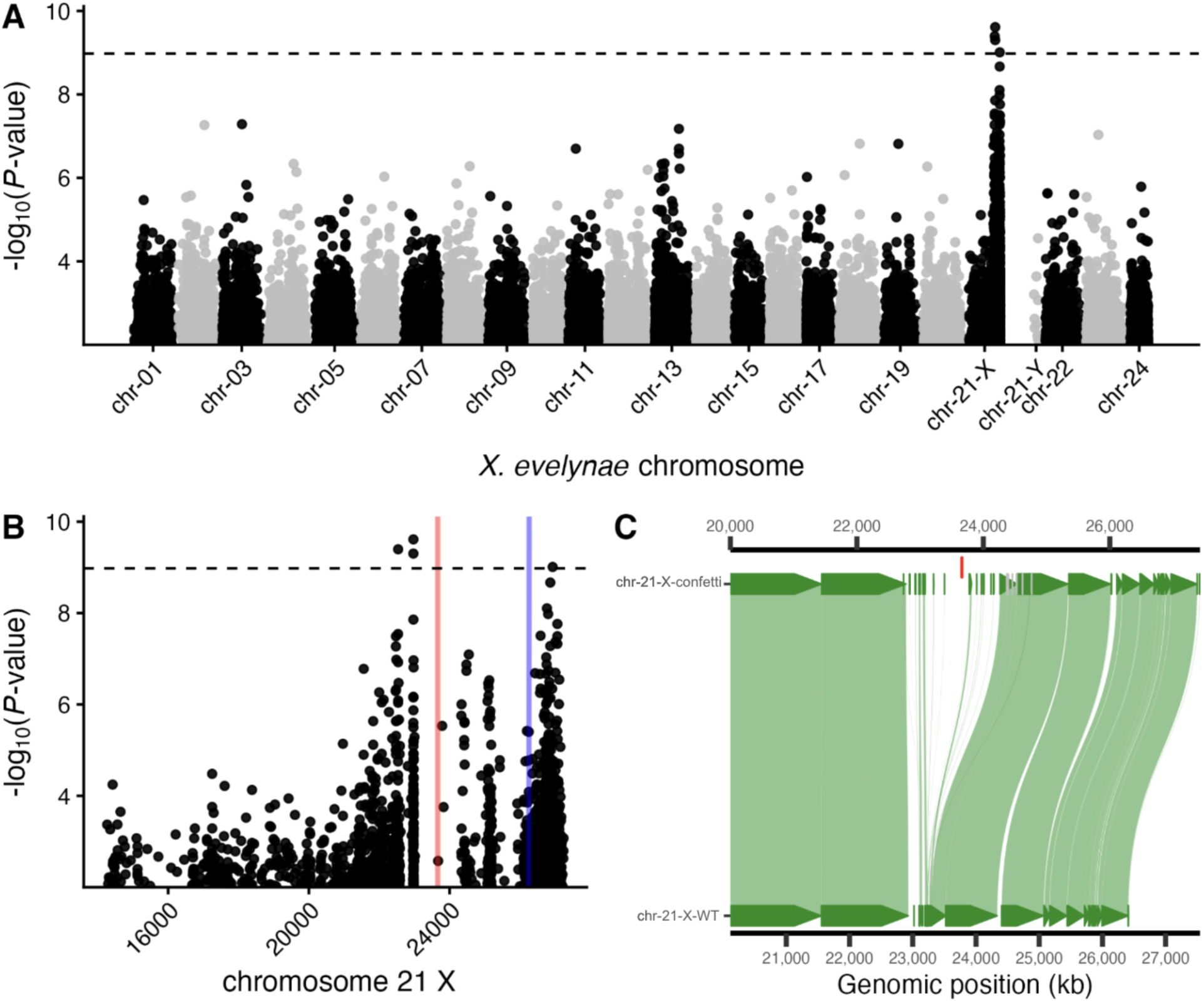
Confetti caudal maps to the genomic region containing *xmrk*. **(A)** Results of a genome-wide association study analyzing allele frequency differences between individuals with confetti caudal (score ≥ 3) and individuals without confetti caudal (score 0). The dashed line indicates the genome-wide significance threshold determined via permutation. **(B)** GWAS results plotted for the distal end of chromosome 21, the X chromosome. The red line marks the location of *xmrk,* the blue line marks the location of *xmrk*’s paralog *egfrb.* **(C)** Chromosome 21 from a long-read assembly of an *X. evelynae* individual with confetti caudal (chr-21-X-confetti) aligned to chromosome 21 from a long-read assembly of an unspotted wildtype individual (chr-21-X-WT). Green areas represent syntenous alignment between the two chromosomes, gray is alignment in the reverse direction (representing inversions). The position of *xmrk* is noted by a segment above the plot in red.

### xmrk is upregulated in pigmented confetti tissue

We performed differential gene expression analysis using bulk RNA sequencing to compare expression across three different tissue types: pigmented caudal fin tissue from individuals with confetti, unpigmented caudal fin tissue from individuals with confetti, and unpigmented caudal fin tissue from individuals without confetti. We were particularly interested in whether any of the genes in the structurally variable region were differentially expressed. We first used PCA to confirm there were no outliers in overall expression patterns of our samples (**Fig. S10**). We found that *xmrk* is significantly upregulated between pigmented caudal fin tissue and tissue from the same region derived from unspotted fish (p < 0.0001, lfc = 5.05; **Fig. 4A, B**). Intriguingly, we also saw that *xmrk* is somewhat upregulated in unpigmented caudal tissue from fish with the confetti pattern compared to unpigmented caudal tissue derived from unspotted fish, indicating a possible role of both genotype and tissue composition (p = 0.015, lfc = 0.580; **Fig. 4B, S11**). Moreover, we also observed a significant difference in expression of *xmrk* between pigmented and unpigmented tissue derived from individuals with confetti caudal (p = 0.011, lfc = 1.01; **Fig. 4B, S11A**). To contextualize these results, we also assessed gene expression of the melanogenesis gene *tyrp1b*, which we found is upregulated only in pigmented tissue, regardless of genotype (**Fig. 4C**), as well as *xmrk’s* paralog, *egfrb,* which has similar expression across all tissue types (**Fig. S12**). We also evaluated whether any other genes within 100 kb of the GWAS interval were differentially expressed by phenotype or tissue type (**Supplementary materials 9**). We found several (n = 10) differentially expressed genes, but none with gene ontology annotations within the melanocyte differentiation or pigmentation GO categories (The Gene Ontology Consortium *et al*. 2026; **Table S2**). We note, however, that four differentially expressed genes had a documented association with pigmentation based on a literature search (**Supplementary materials 9**). We also performed weighted gene co-expression network analysis to identify clusters of co-expressed genes associated with the confetti phenotype, and found that some predicted functional partners of *xmrk,* but not *xmrk* itself, were in gene modules associated with the confetti phenotype (**Table S3**).

**Figure 4:**
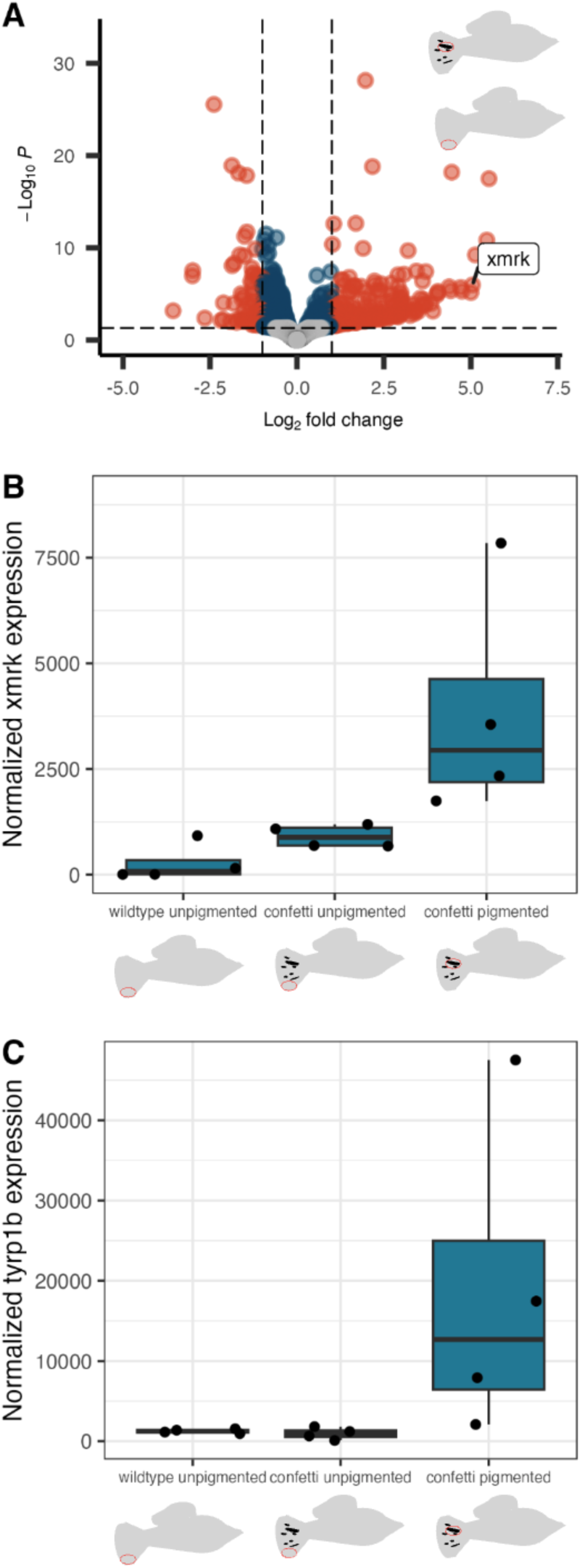
***xmrk* is broadly upregulated in confetti fin tissue (A)** Results of differential expression analysis comparing gene expression in pigmented tissue from individuals with confetti caudal to caudal fin tissue from a wildtype, unspotted fish. Red points indicate genes that cross the significance threshold with an adjusted p-value < 0.05, and also have a log_2_ fold change with magnitude greater than 1. Blue points have p-value <0.05 but have log_2_ fold change < 1. Grey points have p-value > 0.05 and log_2_ fold change < 1. *xmrk* is significantly differentially expressed between these groups (p < 0.0001). **(B)** Normalized expression of *xmrk* across the 3 different tissue types: wildtype unpigmented caudal fin tissue, unpigmented tissue from a confetti individual, and pigmented caudal fin tissue. Each dot is an individual biological replicate. **(C)** Normalized expression of a known melanophore-associated gene (*tyrp1b*) across tissue types.

### *xmrk* does not show evidence of introgression between *X. evelynae* and *X. birchmanni*

One possible mechanism for a repeated role of *xmrk* driving spotting patterns in distantly related species is introgression of this genomic region between *X. evelynae* and the lineage leading to *X. birchmanni* and *X. cortezi*. To investigate this possibility, we compared genetic divergence between the *X. birchmanni* and *X. evelynae xmrk* sequences to the genome-wide background. We found that the observed pairwise sequence divergence between the *xmrk* sequences found in the two species (1.6%) is typical of the genome-wide sequence divergence between them (median 1.5% in 10 kb windows genome-wide; 95% CI: 0.6% - 2.2%). Another possible mechanism of molecular convergence could be incomplete lineage sorting of the causal variants. While we considered this unlikely *a priori* given the deep evolutionary divergence between these lineages, we explored this possibility using simulations implemented in SLiM (**Supplementary materials 10**). We modeled polymorphic variants in an ancestral lineage at a range of frequencies and modeled the inferred divergence time and population sizes of each lineage. We found that, under neutral conditions, maintenance of a shared polymorphism over these timescales is extremely unlikely regardless of the starting frequency; indeed we did not observe this pattern in any of our 2,500 simulations (**Supplementary materials 10**).

### Confetti caudal can develop into melanoma in *X. evelynae* and its hybrids

Several *X. evelynae* individuals with the confetti phenotype were observed to have expanded macromelanophore spots, which appeared to be invading the musculature of the caudal peduncle (**Fig. 5**). We used histology to assess if these expanded spots showed signs of progression to malignancy. We observed an increased number of melanophores in the dermis and macromelanophores invading the fascia of the muscle (**Fig. 5**). According to the *Xiphophorus* melanoma classification scheme used by Schartl *et al*. (1995), this aligns with a stage III melanoma. This is notable since melanomas are more typically found in interspecific hybrid *Xiphophorus*, suggesting that the *xmrk* found in *X. evelynae* may be a particularly potent tumor driver. Intriguingly, we also found potential misregulation of macromelanophore patterning and timing in *X. malinche* × *X. evelynae* F1 hybrids (**Fig. S13**). We found that juvenile F1 hybrids between *X. evelynae* and *X. malinche* developed unusual spotting patterns. Among juveniles phenotyped at 58 days old, many individuals had large macromelanophore spots on their caudal peduncle (**Fig. S13A, B**), a phenotype very rarely seen in *X. evelynae.* In addition to the unusual location of macromelanophores (on the caudal peduncle instead of the caudal fin), F1 hybrids had substantially larger macromelanophore spots than age-matched non-hybrid *X. evelynae* (**Fig. S13**) and spots appeared to develop earlier in hybrids. While some *X. evelynae* have started developing confetti caudal by age 58 days (**Fig. 2A**), they have only one or two very small spots typically not visible without magnification (**Fig. S13C, D**).

**Figure 5:**
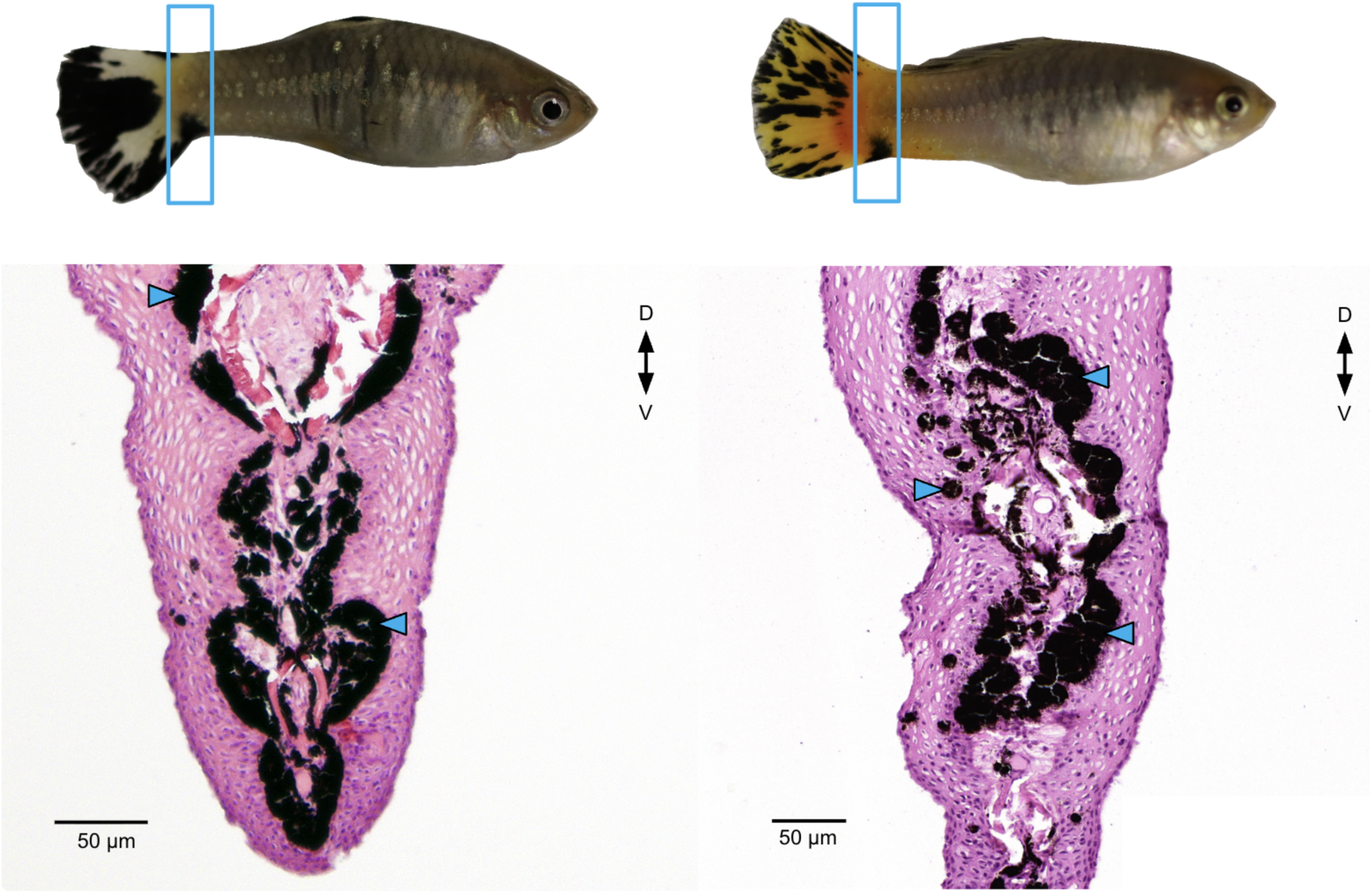
Histology of expanded confetti caudal spots. Top: *X. evelynae* males with confetti caudal pigmentation with an expansion of melanophores into the lower caudal peduncle. The region used for histological analysis is highlighted in the blue rectangle. Bottom: associated cross sections of the caudal peduncle showing macromelanophores invading the muscle fascia (blue arrows). These samples were prepared for histology and stained with a standard Hematoxylin and Eosin stain.

## Discussion

Whether or not evolution is predictable at the molecular level is a foundational question in evolutionary biology. While there is abundant evidence of convergent evolution at the phenotypic level across broad scales of biological diversity, it is still unclear when and under what scenarios we expect to see this convergence mirrored at the molecular level. Cases of convergent phenotypic evolution can help us begin to answer this question. Here, we investigated a convergent macromelanophore tailspot trait in *Xiphophorus* fishes. We found evidence that *xmrk*, a known oncogene and pigmentation gene underlying tail spots in another swordtail species (*X. birchmanni*; Powell *et al*. 2020), is also associated with a trait we call “confetti caudal” in *X. evelynae*. This recurrent use of *xmrk* to encode similar phenotypes allows us to ask key questions about the repeatability of evolution, and begin to provide insight into the evolutionary pressures and ecological forces driving recurrent patterns of pigmentation in *Xiphophorus* species.

The confetti phenotype is polymorphic, present in 66% of wild caught individuals, and more common in females (**Fig. 1D**). Interestingly, while a smaller proportion of males have confetti, they are overrepresented among the most pigmented individuals (11 out of 15 of the individuals in the most pigmented category; Fisher’s exact test: p = 0.000138, **Fig. S14**). This may suggest differences in regulation of the trait between sexes, or may simply reflect differences in the average age of males vs. females collected from the wild, since spots expand with age (**Fig. 2B**). In experiments tracking the development of confetti caudal, macromelanophores became visible at a mean age of 80.6 days (**Fig. 2A**), or just before the expected age of sexual maturation in *Xiphophorus* (Borowsky 1973). However, we also observed cases where individuals developed confetti spots after sexual maturity (**Fig. 2B**). This demonstrates that fish can develop confetti caudal well into adulthood and adds a layer of complexity to our investigations of the genetic basis of the phenotype.

We sought to identify the genetic basis of the confetti caudal trait in *X. evelynae* using a genome-wide association framework. Our GWAS identified one genome-wide significant peak on chromosome 21. This peak was 4.4 Mb wide and overlapped over 100 genes, including *xmrk* (**Fig. 3B**). The peak contains a 1 Mb insertion/deletion structural variant that contains *xmrk* and 5 other genes (**Fig. 3C**). Our GWAS results have a gap in this region, with few reads mapping near *xmrk,* likely due to the segregating deletion, poor read mapping in this repetitive region, or both. To further evaluate this, we relaxed our coverage filtering threshold, and found that a peak centered on *xmrk* is visible using this relaxed threshold (**Fig. S15**). Thus, our results suggest that *xmrk* is associated with the confetti trait in *X. evelynae,* but we cannot yet rule out that another closely linked gene is instead responsible for this phenotype. The *xmrk* gene has long been known to be associated with some macromelanophore pigmentation traits in *Xiphophorus* (Culumber 2014), but genome-wide associations have only been performed in three of the 11 species with macromelanophores (Weis & Schartl 1998) to date (Given *et al*. 2026; Lu *et al*. 2020; Powell *et al*. 2020).

Given evidence of a large structural variant on the putative X chromosome, we wanted to evaluate the proximity of this variant to the sex determining region. We performed a GWAS for sex and found that the sex-determining region colocalized with the GWAS peak for confetti caudal (**Fig. S6**) and with *xmrk*, as in other species (Given *et al*. 2026). Because we were unable to obtain finer resolution through GWAS analysis due to structural complexity in the region (**Fig. S16, S17**), we turned to RNA sequencing to assess the expression of *xmrk* and other genes under the GWAS peak. We found that *xmrk* is strongly differentially expressed between pigmented confetti tissue and tissue from unspotted fish, providing a complementary line of evidence linking *xmrk* to the confetti trait (**Fig. 4A**). Intriguingly, *xmrk* appears to be more highly expressed in unspotted tissue from confetti caudal fish (compared to wild-type fish; **Fig. 4B, S11**). Based on this observation, we reasoned that *xmrk* expression across the whole caudal fin may be consistent with the observation that the confetti phenotype continues to develop and expand over time, sometimes resulting in a completely melanized caudal fin. We also evaluated expression of genes within the GWAS region or within 100 kb on either side (140 genes), and found that a total of 10 other genes were differentially expressed between caudal fin tissue from spotted and unspotted fish. Three of these genes fall into the region encompassed by the structural variant. However, none of the 10 genes fall into annotated pigmentation gene ontology categories.

The finding that *xmrk* is associated with confetti caudal highlights the possibility that the confetti phenotype may be linked to higher risk of melanoma (Walter & Kazianis 2001; Wellbrock *et al*. 2002). In *Xiphophorus,* melanoma originating from macromelanophore pigmentation traits is common in interspecific hybrids but is rarely seen in pure species. *X. variatus* individuals with a macromelanophore body spotting pattern have been shown to occasionally develop melanoma when they reach old age, and spontaneous melanoma formation has also been observed in *X. cortezi* with spotted caudal (Schartl *et al*. 1995). In *X. evelynae*, we found evidence that spontaneous melanoma forms from confetti caudal. Histological analysis of individuals with spots expanding into the caudal peduncle indicated that these expansions show patterns of cell invasion and growth characteristic of malignant melanoma (**Fig. 5**). The findings that spot size continues to grow well into adulthood (**Fig. 2B**) may make it more likely that some melanophore spots continue to expand beyond the caudal fin and invade into the musculature as the fish age. However, the impact of confetti pigmentation on fitness in natural populations will depend on the frequency and age at which melanoma develops from this pattern, and the degree to which it impairs survival (e.g., through swimming performance; Powell *et al*. 2020). If spontaneous melanoma only occurs in older *X. evelynae*, it may be under weak selection in the wild. The impacts of confetti caudal on fitness will not depend on its contribution to melanoma risk alone. While the repeated convergent evolution of macromelanophore tailspots in *Xiphophorus* suggests the trait has an adaptive function, its impacts on fitness are poorly understood. Tailspots have been shown to be sexually selected in *X. cortezi*, but not *X. birchmanni* (Fernandez & Morris 2008; Rosenthal & De León 2006). More broadly, macromelanophore traits have also been associated with variation in aggression and boldness in some *Xiphophorus* species (Fernandez 2010), and based on inferences in other fish, the trait could also confer advantages for shoaling or predator avoidance (Nursall 1973). Future work could use behavior trials and ecological observations to investigate the fitness impacts of confetti caudal in *X. evelynae*.

It is also possible that confetti may be involved in contributing to genetic incompatibilities in hybrids formed with other *Xiphophorus* species, as has been found in the case of other *xmrk-*regulated tailspots in *Xiphophorus* (Lu *et al*. 2020; Powell *et al*. 2020). *X. evelynae* is sympatric with a diverged lineage of *X. malinche* at Juntas Chicas (Aguillon *et al*. 2025), a species that lacks *xmrk* (Powell *et al*. 2020). As a first step toward investigating this possibility, we generated F1 hybrids between this lineage of *X. malinche* and *X. evelynae* with confetti caudal (**Fig. S13**). In other *Xiphophorus* hybrid crosses studied to date, F1 hybrids develop expanded spot phenotypes, while F2 hybrids develop malignant melanoma (Lu *et al*. 2020). Our initial cross results identified a subset of F1 hybrids that developed a large, early-onset macromelanophore spot in their caudal peduncle, a phenotype rarely observed in the parental species (**Fig. S13A, B**). This unique hybrid spot formation suggests there may be misregulation of the confetti trait in hybrids, raising exciting directions for future work on hybrid melanoma in this cross.

We find that, like the similar “spotted caudal” phenotype in *X. birchmanni* and *X. cortezi,* confetti caudal in *X. evelynae* is associated with *xmrk.* Given the evolutionary divergence between these species (**Fig. 1A**), and the fact that *xmrk* arose in the common ancestor via a duplication of the epidermal growth factor receptor B gene (*egfrb*; Du *et al*. 2024; Scarpino *et al*. 2013; Schartl 2008; Weis & Schartl 1998), this suggests that *xmrk* has been repeatedly co-opted for tailspot patterning. While the presence of the *xmrk* gene correlates with the presence of a number of macromelanophore pigmentation traits found in the genus, which occur in different regions of the body (Fernandez & Bowser 2010; Fernandez & Morris 2008; Given *et al*. 2026; Lu *et al*. 2020; Powell *et al*. 2020), association between *xmrk* and macromelanophore traits is complex. The relationship between *xmrk* presence and spotting patterns has only been directly evaluated in a handful of species, and not all species with macromelanophore spotting patterns have the *xmrk* gene (Weis & Schartl 1998). This study highlights a case where the same gene has been repeatedly co-opted to regulate tail spots and presents an exciting first step towards understanding the evolutionary history of spotting traits in *Xiphophorus* more broadly. Despite this convergence in underlying genetic architecture, evidence suggests that the mechanisms by which *xmrk* controls the phenotype are not the same across lineages. In *X. birchmanni,* previous results linked *xmrk* and its upregulation to the spotted caudal phenotype (Powell *et al*. 2020), similar to our findings in *X. evelynae,* but in *X. cortezi,* the association between spotted caudal and *xmrk* appears to be more complex. While *xmrk* is upregulated in spotted tissue in this species (Fernandez & Bowser 2010; Fernandez & Morris 2008), pedigree crosses have shown that the spotted caudal locus in *X. cortezi* has Mendelian inheritance and is autosomally controlled, indicative of a distinct genetic basis (Kallman 1972).

The association between *xmrk*, tail spots, and melanoma also varies between these species. Melanoma formation in non-hybrid *X. birchmanni* is vanishingly rare in the wild (Culumber 2014; Powell *et al*. 2020). By contrast, in *X. cortezi*, melanoma has been observed in young, otherwise healthy fish in both the lab and wild, emanating from the spotted caudal (Fernandez & Bowser 2010; Fernandez & Morris 2008). *X. evelynae* appears to fall in an intermediate category, with spontaneous melanoma forming in the lab in fish of intermediate age, but with less aggressive phenotypes than reported in *X. cortezi* (Fernandez & Bowser 2010; Fernandez & Morris 2008; Schartl et al. 1995). In other words, although the spotting phenotype has a shared genetic architecture between *X. evelynae* and *X. birchmanni*, the incidence of spontaneous melanoma more closely mirrors observations in *X. cortezi*, where the phenotype is autosomally controlled.

## Conclusion

Disentangling the evolutionary history of *xmrk* and its role in macromelanophore traits across species will help us better understand the potential evolutionary paths through which it drives pattern variation. Our findings highlight that convergent evolution at the molecular level can be complex, even when similar traits are associated with the same gene. Moreover, understanding when and how convergent traits emerge through modifications of the same genetic pathways, like those found here, can provide insight into the drivers and constraints of phenotypic evolution and predictability at the molecular level more broadly.

## Acknowledgements

We thank the Federal Government of Mexico and Benemérita Universidad Autónoma de Puebla for permission to collect fish and assistance with permits. We are grateful to the CICHAZ field station for their support of this project. We thank Cheyenne Payne for providing guidance on the analysis of RNA sequencing data. We thank members of the Schumer lab for helpful comments on experiments and/or earlier versions of this manuscript. Stanford University and the Stanford Research Computing Center provided computational support for this project.

## Data Availability

All data for this study will be made fully available without restriction. Phenotype data, images, and files necessary to reproduce figures in the manuscript will be made available on the Dryad Digital Repository. Scripts used for this project will be made available on GitHub.

## Ethics statement

Permission to sample fish was provided by the Mexican government (Permit No. PPF/DGOPA-002/19). Fish were maintained followed procedures described in Stanford Laboratory Animal Care protocol #33071.

## Author contributions

Designed research: S.H.C, S.M.A, M.Schumer

Performed research: S.H.C, N.B.H, J.J.B, T.R.G, T.O.D, K.E.H, A.E.D, A.R, A.P, D.L.P, P.F.Z, M.Schartl, S.M.A, M.Schumer

Analysed data: S.H.C, T.O.D, K.E.H, M.Schartl, S.M.A, M.Schumer

Wrote the paper: S.H.C, S.M.A, M.Schumer

