## Supplemental materials for "Phenotypic and molecular convergence of a cancer-associated pigmentation trait in *Xiphophorus evelynae*"

**Supplementary materials 1:** ImageJ and recolorize analysis to track change in confetti caudal phenotype over time

We analyzed photographs from an experiment where we imaged adults with confetti caudal over a period of several months. We quantified the change in percent melanophore coverage of the fin over time using ImageJ (Schindelin *et al.* 2012) and the R package recolorize (Weller *et al.* 2024). We used ImageJ to extract the tail fin from each photo, and colored the background pink as a contrasting color. We used the recolorize defaults to create an 'initial fit', and then performed the clustered step with a cutoff of 3 (**Fig. S2**). The color mask output for each color was saved as a binary file. We used the mask of the background color to determine the area of the caudal fin by summing the number of pixels in the background mask and subtracting from the total pixels present in the image. The masks for black color were also summed to quantify the number of pixels in black confetti spots. We visually checked the masks for accuracy. In some cases, multiple masks needed to be added to capture the confetti melanized area, given variation in lighting conditions across photographs. For these individuals, several masks were summed to calculate the total pixels present in melanized confetti caudal spots. To determine the fraction of the caudal fin covered in melanized confetti caudal spots, we divided the sum of the melanized pixels by the number of pixels in the entire tail. Fish that developed confetti caudal after tracking began were considered to have 0 melanophore coverage at timepoints before phenotype onset. Results were plotted in R version 4.1.0 using the tidyverse package (Wickham *et al.* 2019).

**Supplementary materials 2:** Comparing quantitative recolorize scoring with qualitative scoring

Our recolorize scoring approach gives quantitative results but is labor intensive and requires uniform lighting conditions. We thus wanted to compare this approach to the qualitative phenotyping scoring used for the rest of the experiments from wild-caught individuals. To do so, we performed recolorize quantification on a subset of the wild-caught photoset used for the GWAS. We selected individuals 1-35 from the dataset, and followed the same procedure described above. Individuals without confetti (scored as 0) were not analyzed with recolorize, and were considered to have a melanized fraction of 0 (n = 8). For some individuals (n = 9) with low confetti scores, the recolorize settings used for the adult tracking experiment failed to clearly differentiate melanophore confetti spots from the background color, likely due to the different lighting conditions of photos taken in the field. To correct this issue without changing the recolorize settings, these photos were edited in FIJI (Schindelin *et al.* 2012). Each confetti spot was manually outlined using the polygon tool and filled in with a contrasting color prior to recolorize analysis. The confetti fraction from this quantitative approach was strongly correlated with the qualitative confetti scoring metric used in the GWAS (Spearman rank correlation = 0.94, $p < 0.0001$), suggesting that our scoring approach accurately reflects phenotypic variation (**Fig. S3**).

**Supplementary materials 3:** Comparison of developmental timing of confetti caudal in *X. evelynae* to spotted caudal in *X. birchmanni*

In order to further assess differences between the confetti caudal phenotype and tailspot phenotypes in related species, we compared the timing of development of confetti caudal with spotted caudal in *X. birchmanni* by phenotyping photographs available from a previously collected developmental time series (Dodge *et al.* 2024). Because the *X. birchmanni* dataset contained only males, we restricted our comparison to males in *X. evelynae* as well. We evaluated patterns in *X. evelynae* (n = 6) and *X. birchmanni* (n = 12) males who had developed tailspots by the final day of the developmental tracking experiment. Exact age information was not available for the previously collected *X. birchmanni* dataset, so we instead used gonopodium development score, a morphological classification of male sexual maturity, as a proxy for male age. We compared the fraction of *X. birchmanni* and *X. evelynae* who had already developed tailspots at gonopodium score one (when they were first able to be identified as males, following the gonopodium score guide in Dodge *et al.* 2024).

As another step to make the datasets more directly comparable, we adjusted our scoring approach for early stage confetti caudal in *X. evelynae*, since the *X. birchmanni* study did not include imaging under the dissecting microscope. As such, we required individuals in the *X. evelynae* dataset to have a confetti score of 1, a phenotype that could be visualized without magnification. Thus, *X. evelynae* individuals with a confetti score of 0.5 (melanophores visible under magnification but not with the naked eye) were converted to score 0, for comparison to *X. birchmanni*. We tested for a difference in the ratio of individuals who had developed tailspots at gonopodium stage 1 using a Chi-squared test ($\chi$-squared = 1.8591, df = 1, p = 0.1727; **Fig. S5**).

**Supplementary materials 4:** Assessing evidence of population substructure

We used a simple scan for allele frequency differences between spotted and unspotted individuals for GWAS. However, this approach can suffer from a high false positive rate in the presence of population structure, particularly when the phenotype is associated with substructure. In order to evaluate whether this is likely to be an issue in our analysis, we performed a PCA and tested for correlations between the major PCs and phenotype in our data. Since our data is low coverage, in order to generate a data frame suitable for PCA, we randomly sampled a read from each position in a pileup file generated with BCFtools (Danecek *et al.* 2021) to generate pseudo-haploid calls for each individual. We assigned the sampled allele as the genotype at that position and converted remaining uncovered sites to NAs. We used the recode function in PLINK (Chang *et al.* 2015) to remove sites with a minor allele frequency below 5% or with missing data in more than 75% of individuals. We used PLINK to perform PCA and plotted the results. We see no strong differentiation between confetti and non-confetti individuals in PC space, suggesting that population sub-structure is unlikely to be driving our GWAS peak. PCs 1-3 explained 4.9%, 4.3%, and 3.8% of the variation in the SNP data respectively. We tested for correlations between PC1 and phenotype (Pearson's $r = 0.12$, $p = 0.11$), PC2 and phenotype (Pearson's $r = 0.05$, $p = 0.50$), and PC3 and phenotype (Pearson's $r = 0.16$, $p = 0.04$) and found no or marginal correlations between these PC loadings and phenotypic variation (**Fig. S8**). We thus proceeded with the assumption that population structure is not driving our GWAS results.

**Supplementary materials 5:** Analysis of the structural variant region using low coverage data from the GWAS population

Comparison of long-read assemblies indicated that the haplotype lacking *xmrk* had a large 1 Mb structural variant relative to the haplotype containing *xmrk.* This led us to hypothesize that the large structural variant may be broadly correlated with trait presence/absence. We estimated coverage in sliding windows using samtools in this region between cases and controls. We filtered alignments with MQ < 30 and normalized coverage per window to the median coverage chromosome wide in cases and controls. We find substantially reduced coverage in the structurally variable region, which colocalizes with our GWAS peak, in unspotted compared to spotted fish (blue versus black in **Fig. S9**), consistent with a role for the structural variant in phenotype presence/absence. We repeated this analysis in a limited number of individuals for which we had both phenotypic information and high coverage PacBio data, focusing on coverage in the *xmrk* region using pbmm and visualization in IGV (see main text). This analysis confirmed both low mappability in both sample sets and the presence of a deletion co-localizing with *xmrk* in unspotted individuals (**Fig. S9**).

**Supplementary materials 6:** Coexpression clustering analysis

In order to further interpret the results of our RNAseq analysis, we performed weighted gene co-expression network analysis (WGCNA) to identify clusters of coexpressed genes associated with the confetti phenotype. This analysis clusters genes into 'modules' with highly correlated expression patterns. We imported the output of our differential gene expression (DGE) analysis conducted with DESeq2 into R to conduct analysis using the package WGCNA, following the approach described in Payne *et al.* 2022. Briefly, the gene list was filtered to remove all genes with zero counts for more than half of the samples or with 0 variance (1308 were removed in this step, leaving 23,447 genes). We chose a soft-thresholding power by calculating the scale-free topology fit index for powers ranging from 1 to 30. We selected the asymptote of soft-thresholding power using the recommended thresholds for scale-free topology fitting index R2 > 0.8 and mean connectivity <100 (Zhang & Horvath 2005). The results of this analysis asymptote at a value of 9-10; based on this result, we selected a soft-thresholding power of 9. We used this soft-thresholding value and the WGCNA blockwiseModules function to construct a single-block unsigned network. We used a maximum block size of 24,000 set to a value larger than the number of genes in our dataset (n = 23,447), and a minimum module size of 20. We used an unsigned topological overlap matrix to cluster genes by strength of coexpression without incorporating expression direction. In order to identify coexpression modules correlated with variation in the confetti trait, we tested for correlations between the module eigengene and either phenotype (confetti vs. non-confetti) or tissue type (melanized vs. not melanized) using the WGCNA corPvalueStudent function. We selected modules that were correlated with spotting phenotype, pigmented tissue, or both at a p-value <0.05. Thirteen modules were significantly correlated with pigmented or wildtype tissue.

We assessed whether *xmrk, egfrb,* and genes identified by a STRING database search as likely interacting genes with *xmrk* were present in any of these modules. We reasoned that this may identify important interactions in the same pathway, as there are likely many genes downstream of *xmrk* required to produce and pattern the macromelanophore cell type of confetti spots. We found that *xmrk* was not present in any significant module. None of the predicted functional partners of *xmrk* were present in any module. *Egfrb* was present in one module (blue), which was differentially expressed in tissue from unspotted fish compared to spotted fish (p = 0.0394), and was positively associated with the wildtype phenotype (module-trait relationship 0.599) (**Table S3**). Despite this, *egfrb* is not differentially expressed between groups in the original analysis (**Fig. S12**).

**Supplementary materials 7:** Generation and phenotyping of *X. malinche* × *X. evelynae* F1 hybrids

Our GWAS indicated association between the region containing *xmrk* and the confetti trait, and we also observed spontaneous melanoma formation in *X. evelynae* individuals. Past work on the role of *xmrk* in pigmentation has repeatedly linked it to the development of hybrid melanomas, including in other species with macromelanophore tailspots. *X. evelynae* in the Juntas Chicas population are sympatric with only one other *Xiphophorus* species, a lineage of *X. malinche*, and the species pair does occasionally hybridize in the wild (Haghani *et al.* 2025). Moreover, *X. malinche* lacks the *xmrk* gene and hybrids between *X. malinche* and another species with *xmrk* have a high risk of developing hybrid melanoma (Powell *et al.* 2020). We generated F1 hybrids to test if hybridization between *X. evelynae* and *X. malinche* was possible, and if F1 hybrids showed evidence of changes in the macromelanophore patterns that may suggest misregulation of *xmrk.* We produced artificial crosses between male *X. evelynae* with confetti and female *X. malinche.*

**Supplementary materials 8:** Reported macromelanophore tail spot patterns in *X. variatus*

In the main text, we discuss *X. evelynae* as being unique among the platyfish clade in having a macromelanophore tailspot. However, a macromelanophore tailspot pattern described as "spotted caudal" has been reported in a population of *X. variatus*, which is a close relative of *X. evelynae.* This spotting pattern is reported to only occur in an *X. variatus* population in the Río Cazones and Río Tuxpan system, in geographical proximity to the Juntas Chicas site (Kallman 1972; Rosen 1960), and has not been observed across the rest of *X. variatus'* wide range. We treat these reports with caution because *X. variatus* and *X. evelynae* are difficult to distinguish phenotypically, and *X. evelynae* was considered a sub-species of *X. variatus* at the time this phenotype was first reported (Rosen 1960). Due to changing species distinctions, we are unsure whether the observed *Xiphophorus* with macromelanophore tail spotting patterns were instead *X. evelynae,* or if some populations of *X. variatus* do possess a distinct spotted caudal-like pattern. Thus, we treat the tail spotting phenotype of *X. variatus* as ambiguous in our discussions in the main text.

**Supplementary materials 9:** Expression of all genes under the GWAS peak

We analyzed evidence for differential expression of all genes under the GWAS peak between groups. We determined the start and end position for the peak on chromosome 21 based on the location of any SNP passing the significance threshold, and added a 100 kb buffer on either side to be conservative. This region contains 140 genes. Of these genes, 103 passed our RNAseq filtering criteria (see main text), and 11 were differentially expressed between pigmented caudal tissue in confetti individuals versus unpigmented tissue from an unspotted fish, including *xmrk* (**Table S2**).

To determine whether any of these differentially expressed genes may be additional candidates for regulating the confetti caudal phenotype, we investigated whether they had known roles in pigmentation. None of the other genes have gene ontology annotations within the melanocyte differentiation or pigmentation GO categories (The Gene Ontology Consortium *et al.* 2026). We then performed Google Scholar searches for the gene name and the terms 'pigmentation' and 'melanophore,' and noted cases where the gene had been previously associated with pigmentation or melanoma in the scientific literature. Genes *eif2s3*, *mc4r*, *b4galt2*, *scn5a*, *rpl22l1*, and *acvr2b* had no documented association with pigmentation. Inhibition of *eif5a* has been found to impair tumor growth in murine melanoma cell lines (Jasiulionis *et al.* 2007). This gene had lower expression in pigmented confetti tissue compared to tissue from an unspotted fish in our analysis. *cdh6*, which may play a role in xanthophore pigmentation polymorphism in *Xiphophorus maculatus* (Soria et al. 2025), is highly expressed in melanomas induced by transgenic ectopic expression of *xmrk* in Medaka (Abdulsahib *et al.* 2024), and mutations of *cdh6* have been found in skin cutaneous melanoma in humans. *cdh6* had higher expression in pigmented confetti tissue compared to tissue from an unspotted fish in our analysis. We note that *cdh6* fell within the 100kb flanking region, but not within the GWAS peak itself. A reduction in *Drosha* protein levels has been observed in human melanoma cell lines (Mione & Bosserhoff 2015). *Drosha* had higher expression in confetti tissue in our analysis. *sat1* is involved in polyamine catabolism, and the polyamine pathway is involved in human skin pigmentation (Sridharan *et al.* 2020). *sat1* had lower expression in unpigmented tissue from a confetti fish compared to tissue from an unspotted fish in our analysis.

**Supplementary materials 10:** Sequence divergence of *xmrk* and SLiM simulations of incomplete lineage sorting

Beyond convergent evolution of tailspot patterns, there are two additional possibilities for why tail spotting traits may be found in distantly diverged species. These include introgression of the causal variants between *X. evelynae* and *X. birchmanni* (or *X. cortezi*), or incomplete lineage sorting resulting in unusual phylogenetic distributions. To evaluate these possibilities, we used two approaches. *X. evelynae* and *X. birchmanni* are deeply diverged, such that we would expect introgression to leave a detectable signature on pairwise sequence divergence (Dxy). We calculated Dxy between *xmrk* copies found in *X. evelynae* and *X. birchmanni*, finding that divergence in this alignment is 1.6%. To compare to the genome-wide background, we used pseudoreferences generated from short-read Illumina data for a previous project (Preising et al. 2024) and calculated Dxy in 10 kb windows between the two species. Divergence between *xmrk* in the two species fell squarely within the genome-wide distribution (see main text).

We next turned to the hypothesis of incomplete lineage sorting. *X. evelynae* and the lineage leading to *X. birchmanni* and *X. cortezi* are deeply diverged, and the pattern of only three species having this trait seems unlikely to be generated by ILS. Specifically, if tail-spot patterns linked to *xmrk* were maintained as polymorphisms in the common ancestor of these two lineages, we may expect to see this pattern more commonly across the 19 daughter lineages. Nevertheless, to directly investigate this possibility, we performed neutral simulations in SLiM intended to quantify the rate of ILS over these timescales. To parameterize the simulations, we used the inferred time-averaged effective population sizes for *X. birchmanni*, *X. cortezi*, and *X. evelynae* from previous work (51500-57500; Powell et al. 2021; Haghani et al. 2025) and calculated the expected divergence time using the relationship: Tdiv/2Ne = Dxy/theta - 1. Measures of Dxy and theta were drawn from previous work. Importantly, estimated theta for each species is comparable, ranging from 0.0008-0.0012 (Powell et al. 2021;

Haghani et al. 2025). These estimates allowed us to infer divergence time between each lineage for the SLiM simulations. We initialized the simulations with a range of starting allele frequencies for the polymorphism in the common ancestor: 0.1, 0.2, 0.3, 0.4, and 0.5 and tracked the maintenance of the polymorphism until fixation in all tracked lineages or the end of the simulations. We performed 500 replicate simulations for each ancestral allele frequency, and found that the polymorphism was not maintained in any simulation over the relevant timescales under these neutral conditions (see main text).

**Supplemental figures**

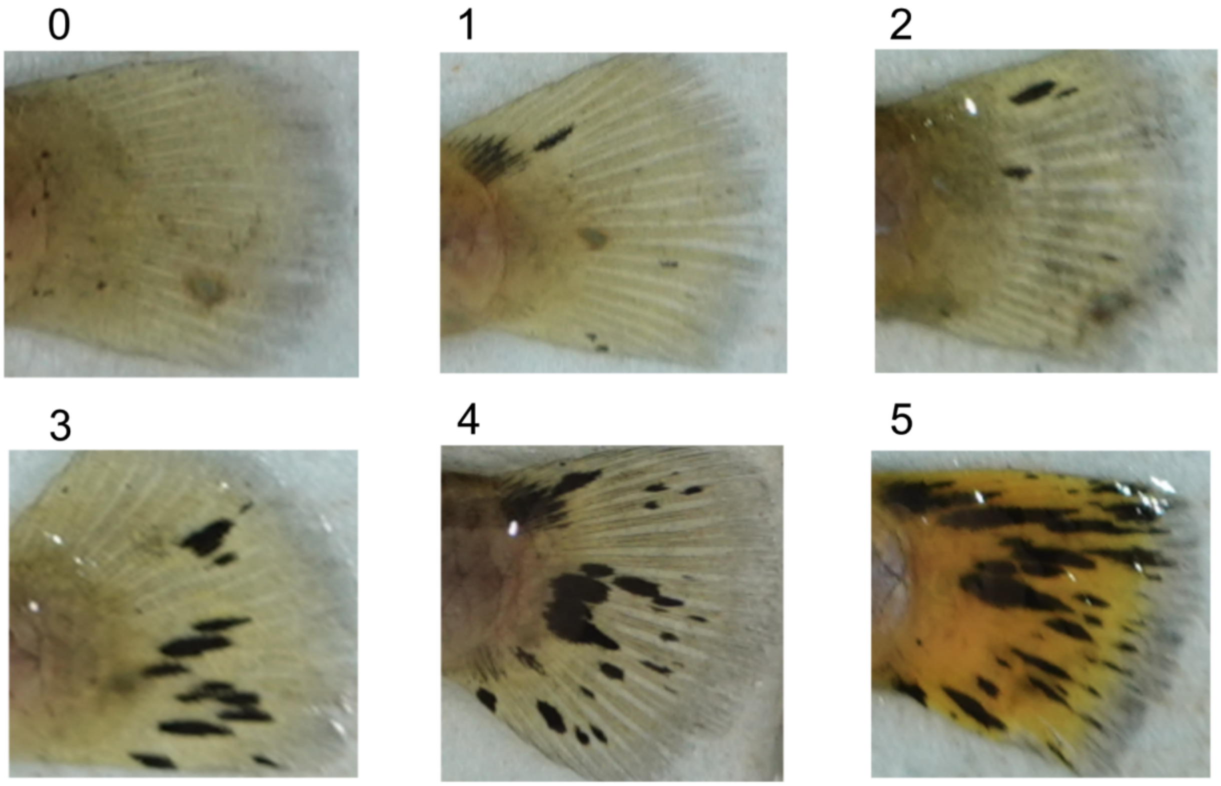


**Figure S1: Phenotypic variation in the confetti caudal trait and associated categorical scoring**
Samples were given a phenotype score ranging from 0, if macromelanophore spots were completely absent, to 5, where approximately 30% of the caudal fin was melanized.

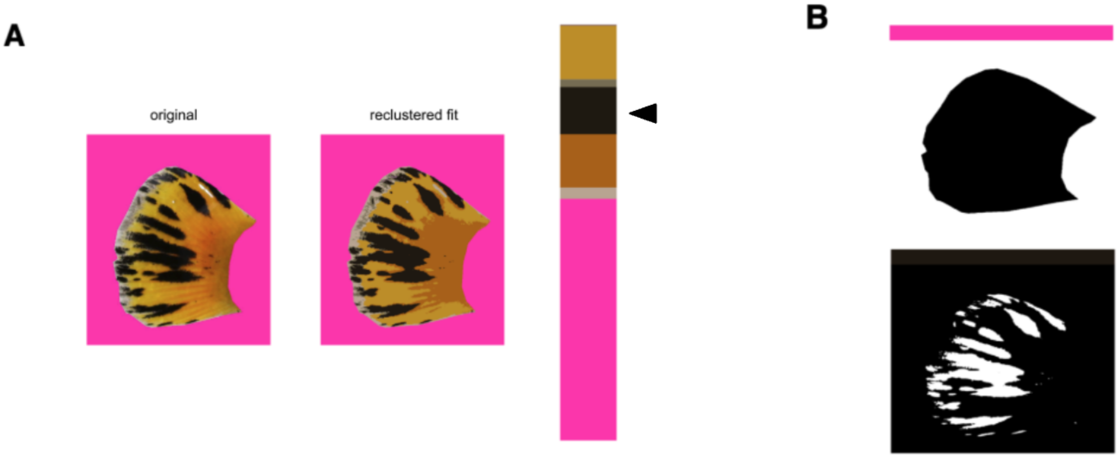

**Figure S2: Quantification of caudal macromelanophore area in confetti individuals using recolorize**

**(A)** Example output from the recolorize workflow, including the original photo with background removed, recolorized reclustered fit, and proportion of pixels in each color category (represented as a single bar). The black color mask (indicated with a black arrow head) was used for confetti caudal area. **(B)** The full tail area mask (top) and black color area mask (bottom) for the individual shown in **A**, were used to calculate the fraction of melanophore coverage for each individual.

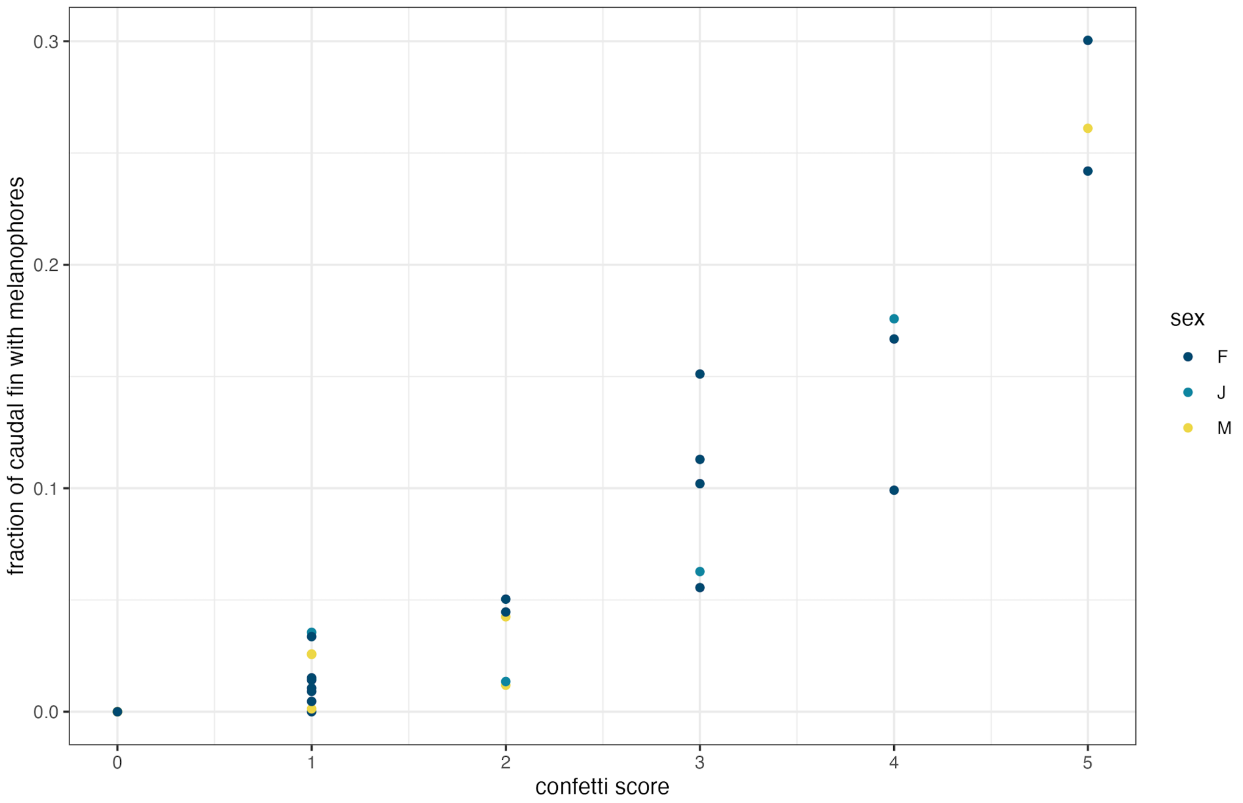


**Figure S3: Comparison of qualitative and quantitative scoring methods**

**(A)** Categorical confetti score for a subset (n = 35) of wild-caught individuals vs fraction of melanophore coverage using recolorize (quantitative score) for the same individuals. These results highlight a strong association between our quantitative and qualitative scoring approaches (Spearman rank correlation = 0.94, $p < 0.0001$).

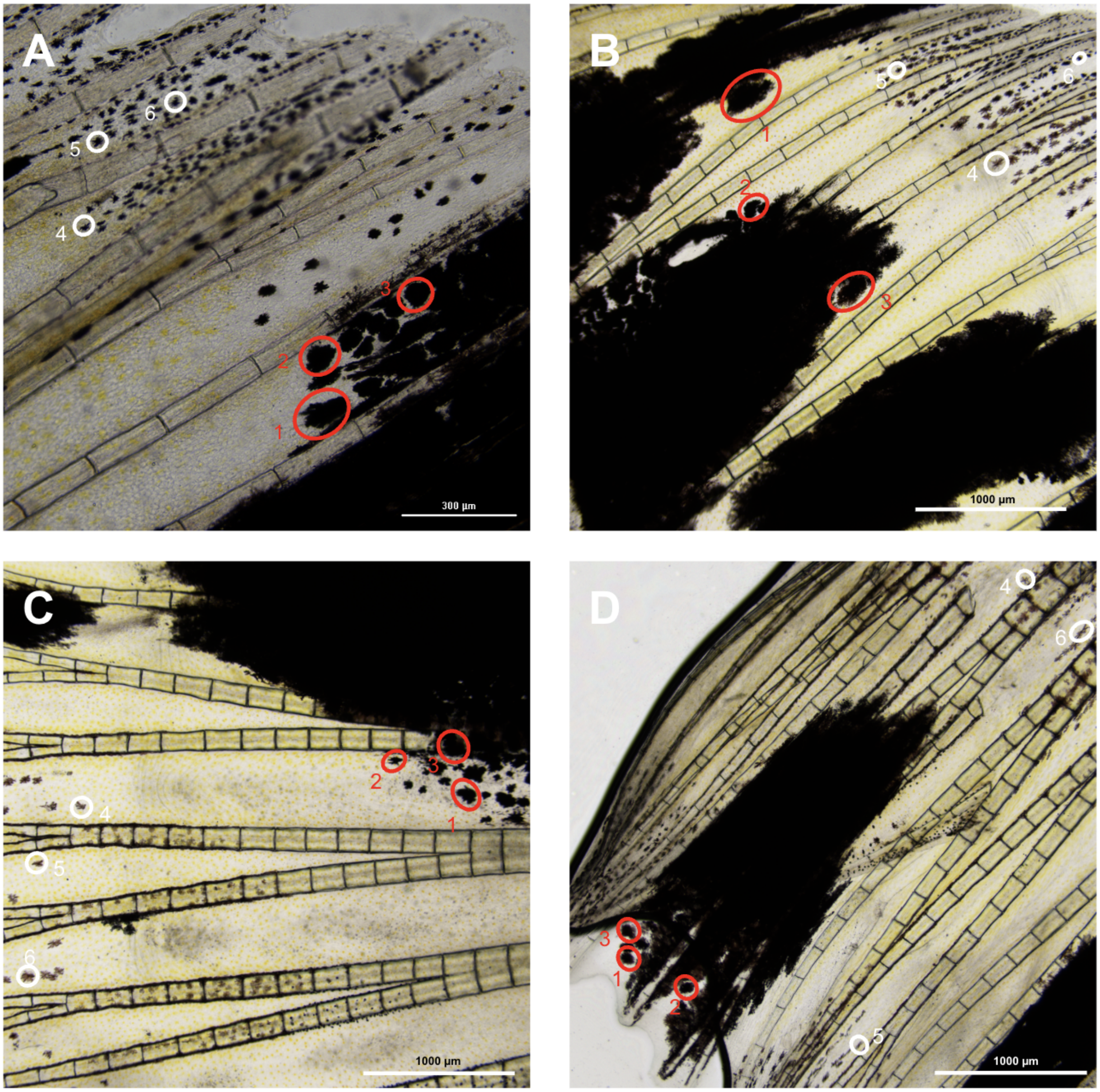


**Figure S4: Images showing pigment cells on the caudal fin under 4X magnification to quantify melanophore sizes**
Melanophores that form part of the confetti pattern are larger than dispersed melanophores on the caudal fin. **(A-D)** Images of the caudal fin of *X. evelynae* individuals with confetti caudal. Circles indicate the melanophores that were measured (Table S2); red circles indicate melanophores that are part of the confetti trait, white circles indicate other melanophores dispersed across the fin that do not contribute to the confetti phenotype.

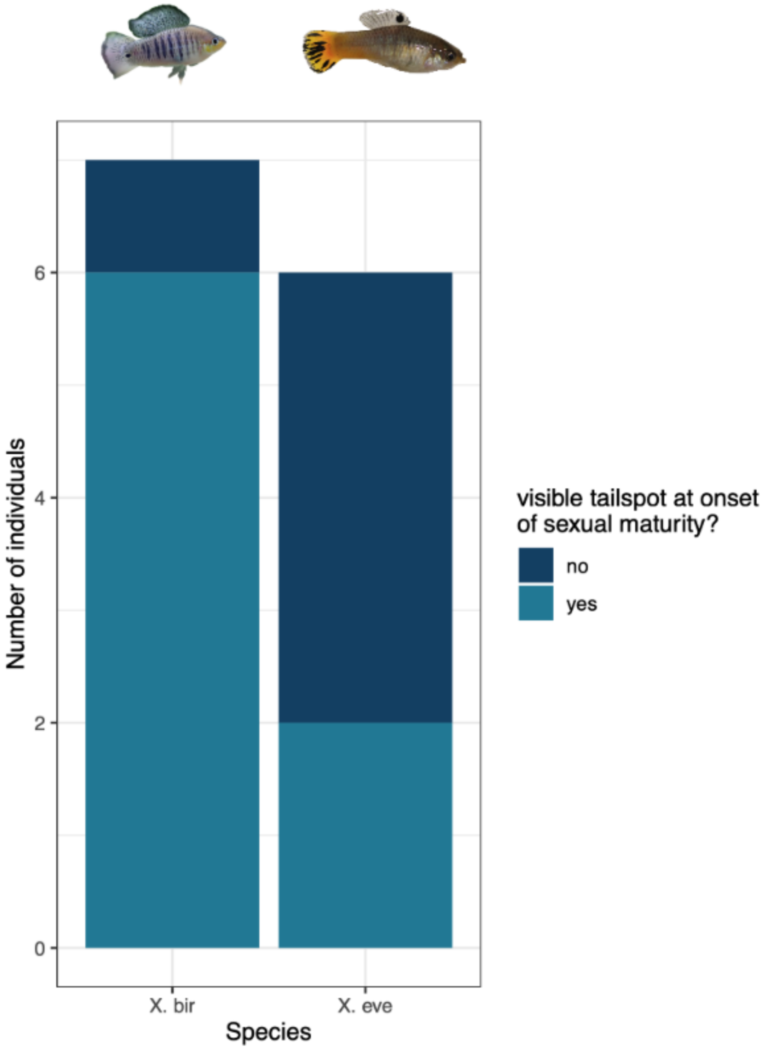


**Figure S5: Comparison of developmental timing of tailspot phenotypes in *X. evelynae* and *X. birchmanni* juvenile males**
See Supplementary materials 4 for details.

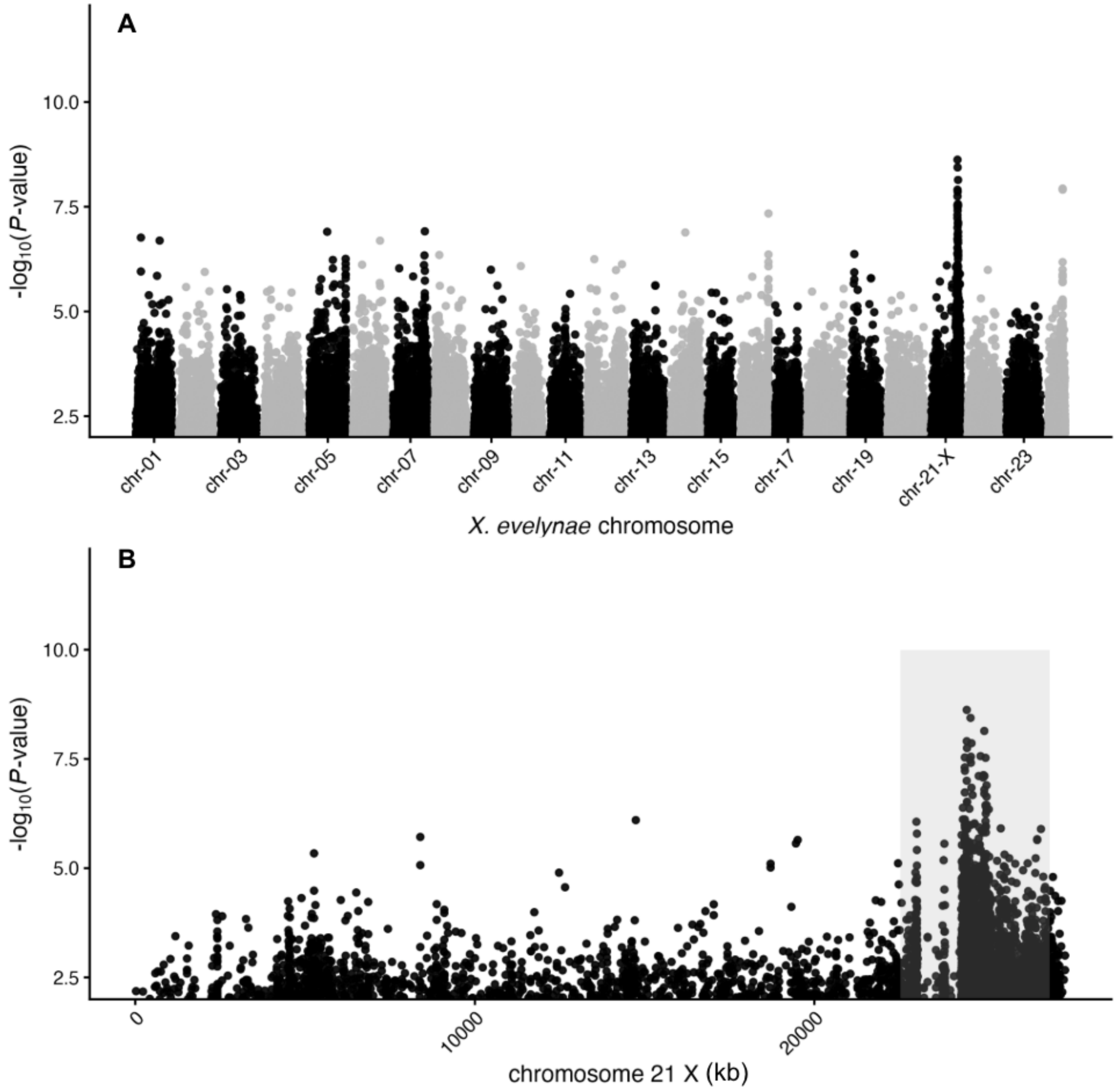


**Figure S6: The sex determining region in *X. evelynae* overlaps with the confetti GWAS peak**

**(A)** Results of a genome-wide association study for allele frequency differences between individuals phenotyped as adult males vs. adult females. The GWAS peak for sex overlaps with the peak for the confetti phenotype. Note that this plot shows GWAS results without coverage filtering. **(B)** A zoomed in view of the distal end of chromosome 21 (the X chromosome). The gray box marks the location of the GWAS peak for the confetti trait.

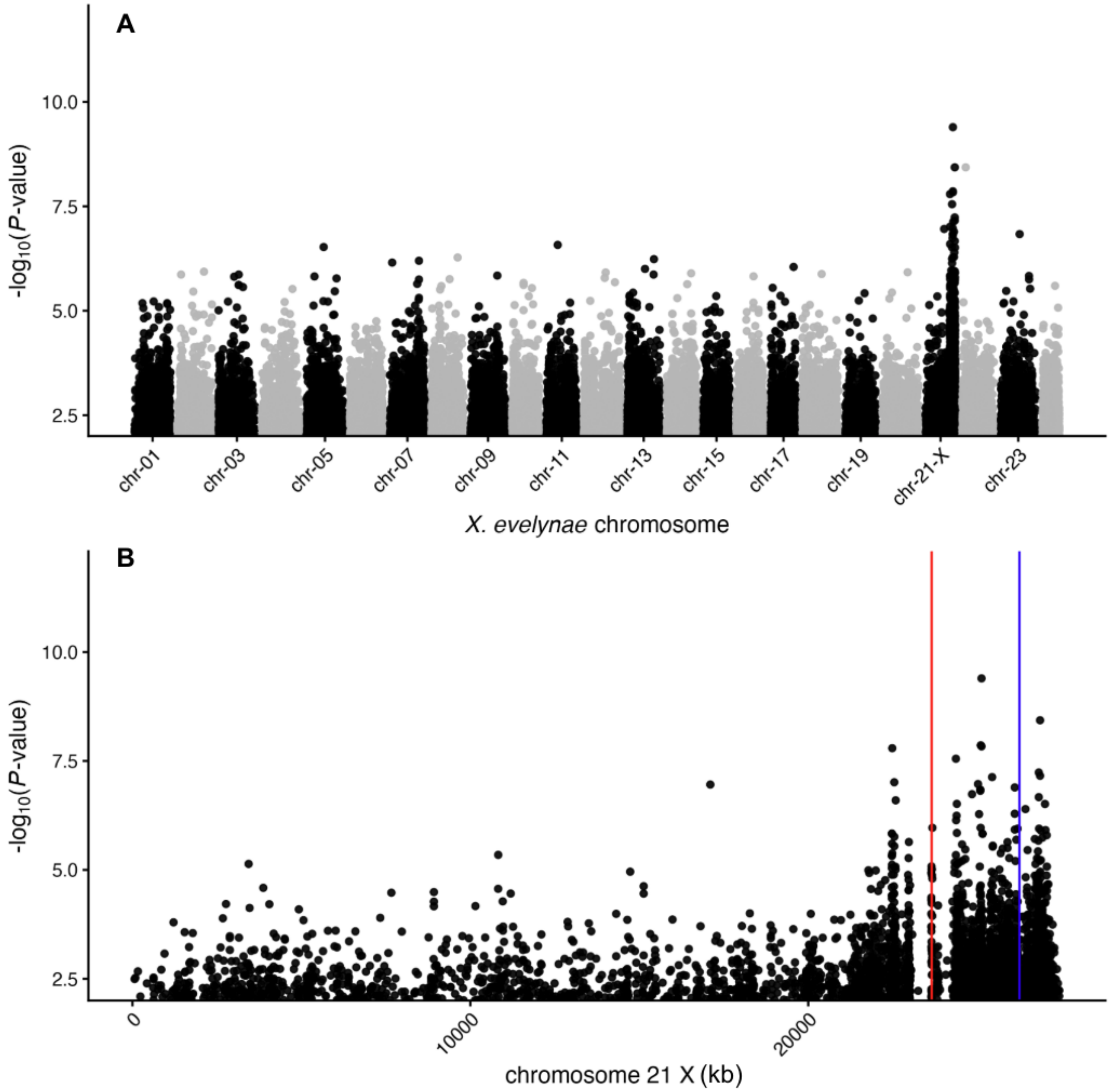


**Figure S7: GWAS for the confetti trait limited to female individuals only**
**(A)** Results of a genome-wide association study for allele frequency differences between individuals with confetti (score ≥3) and individuals without confetti (score 0), limiting only to the adult females in our dataset (n = 75). This analysis was performed excluding the Y chromosome. Note that this plot shows raw results without coverage filtering. **(B)** A zoomed in view of the distal end of chromosome 21 (the X chromosome). The red line marks the location of *xmrk,* the blue line marks the location of its paralog *egfrb.*

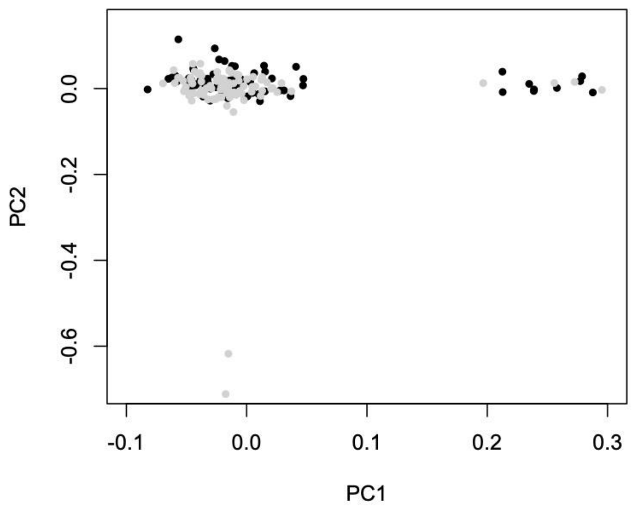


**Figure S8: Population substructure does not correlate with phenotype in our GWAS dataset.** Plotted here are PC1 (4.9% of SNP variation) and PC2 (4.3% of SNP variation). Dark dots indicate individuals with the confetti phenotype and gray dots indicate unspotted individuals. The PCA suggests no clustering based on phenotype.

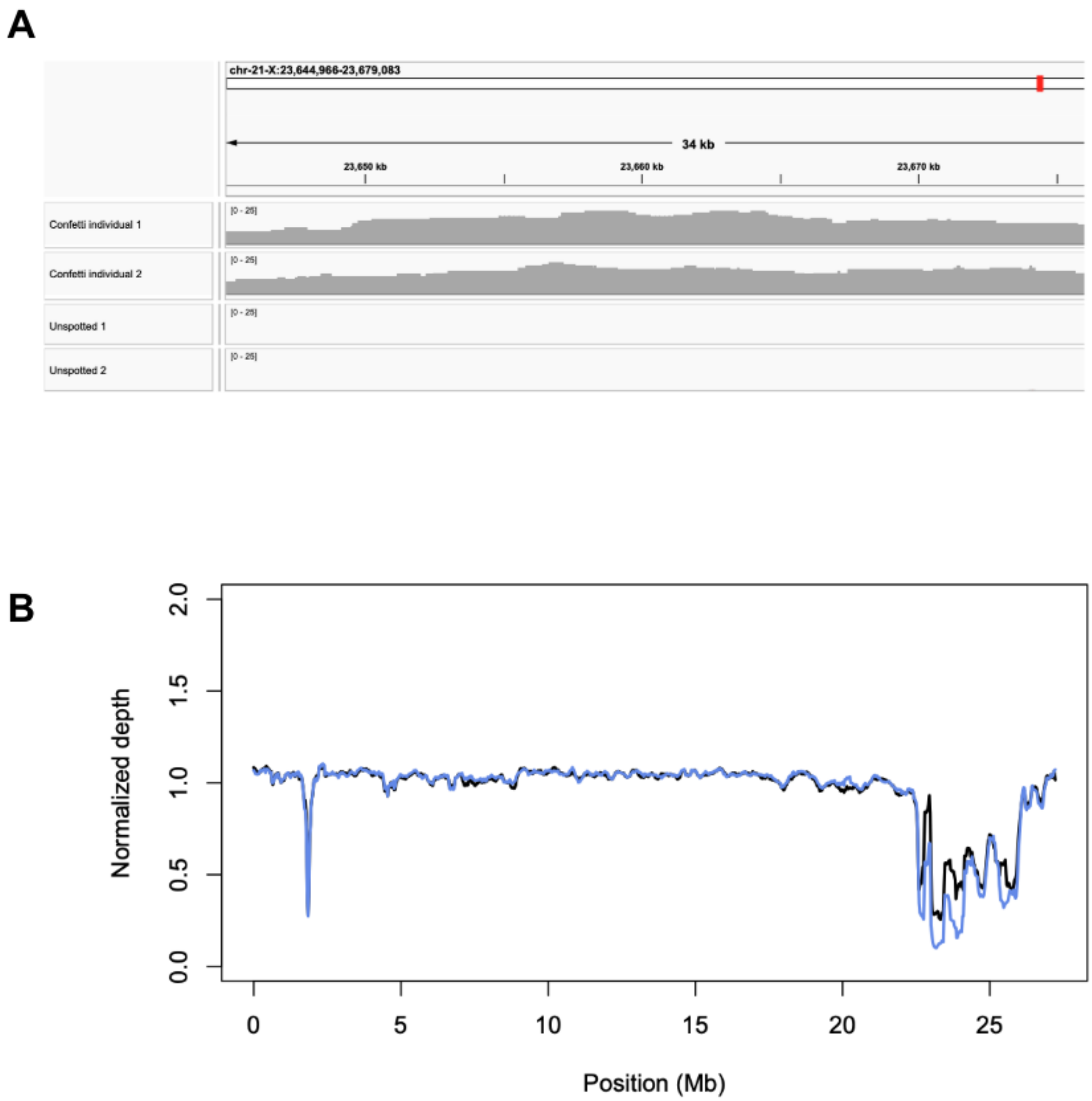


**Figure S9: Reduced coverage in the structurally variant region in WT compared to confetti fish**
**(A)** Coverage results for PacBio data mapped to the *X. evelynae* reference genome without the 1 Mb presence/absence variant highlights differences between the two spotted and two unspotted individuals. Results shown here were visualized in IGV and display the *xmrk* region. **(B)** Estimated coverage depth in sliding windows of chromosome 21 (the X chromosome). Blue line indicates wildtype unspotted individuals, black line indicates individuals with the confetti phenotype. There is substantially reduced coverage in the region of the GWAS peak between wildtype and confetti individuals.

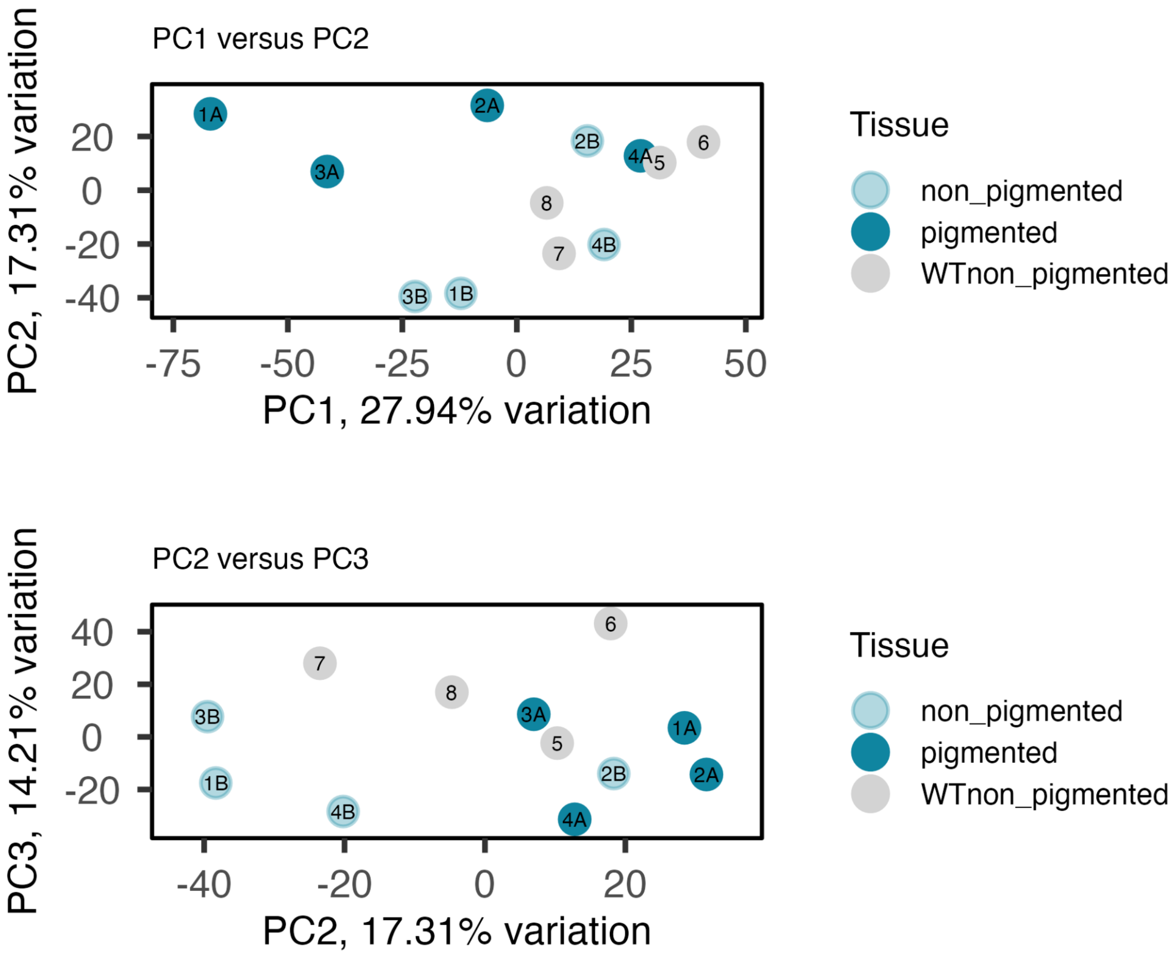


**Figure S10: Overall patterns of gene expression of confetti and wildtype individuals in PC space**

Gene expression of caudal fin tissue from unspotted wildtype fish, pigmented tissue from confetti fish, and unpigmented tissue from confetti fish. Top: PC1 vs PC2. Bottom: PC2 vs PC3. No outliers were identified, thus all samples were retained in downstream analyses.

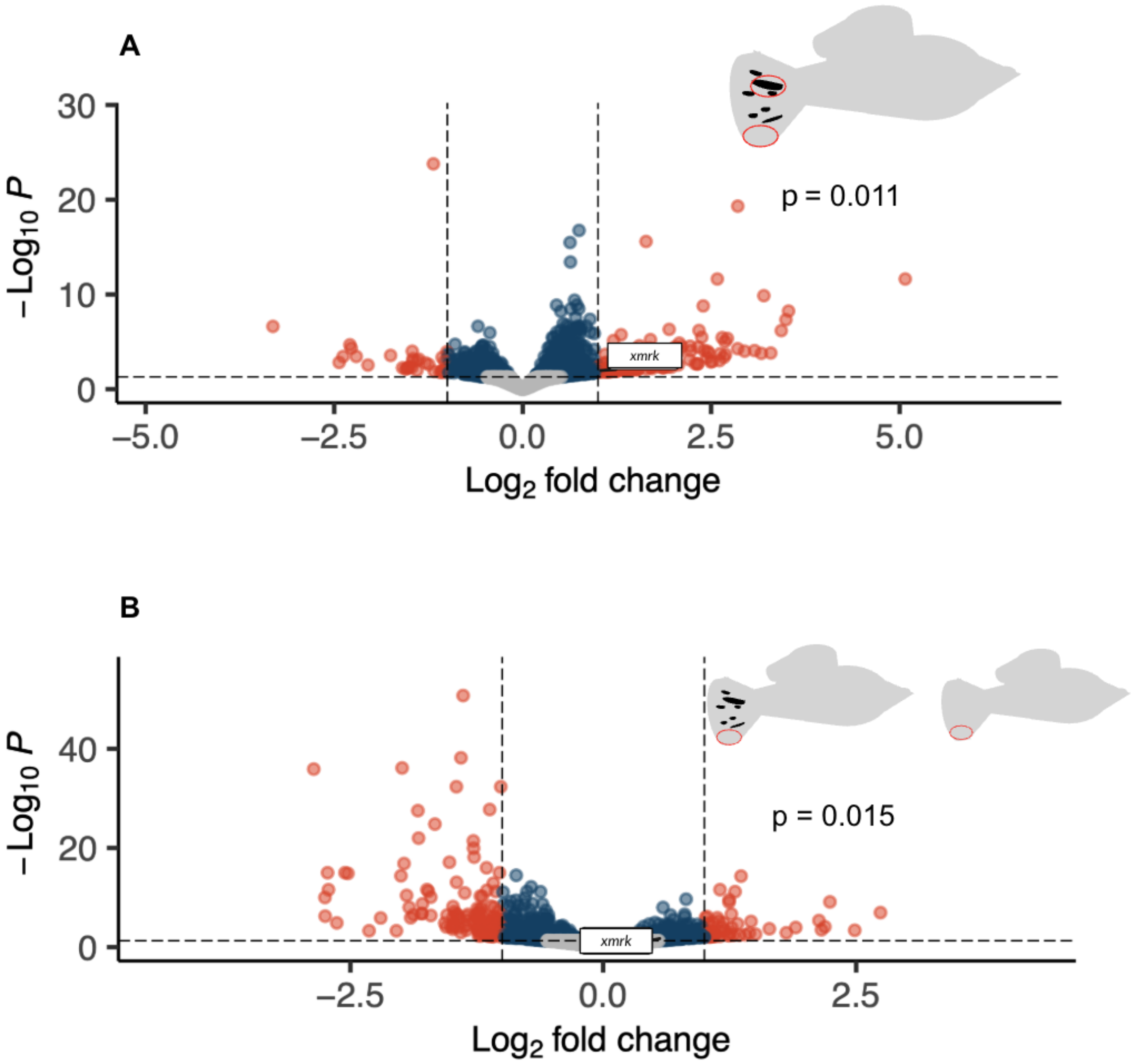


**Figure S11: Additional gene expression analysis and comparisons of *xmrk***
**(A)** Results of differential gene expression analysis comparing gene expression in pigmented tissue from confetti individuals to paired unpigmented tissue from the same individuals (comparison highlighted in inset images). Red points indicate genes that cross the significance threshold of adjusted p-value < 0.05 and also have a $log_2$ fold change with magnitude greater than 1. Blue points have p-value <0.05 but have $log_2$ fold change < 1. Grey points have p-value > 0.05 and $log_2$ fold change < 1. *xmrk* is differentially expressed in this comparison with a modest $log_2$ fold change (lfc) (p = 0.011, lfc = 1.01). **(B)** Results of differential gene expression analysis comparing gene expression in unpigmented tissue from confetti individuals to unpigmented tissue from wildtype fish (comparison highlighted in inset images). *xmrk* is differentially expressed (p = 0.015, lfc = 0.580) in this analysis but with a modest $log_2$ fold change.

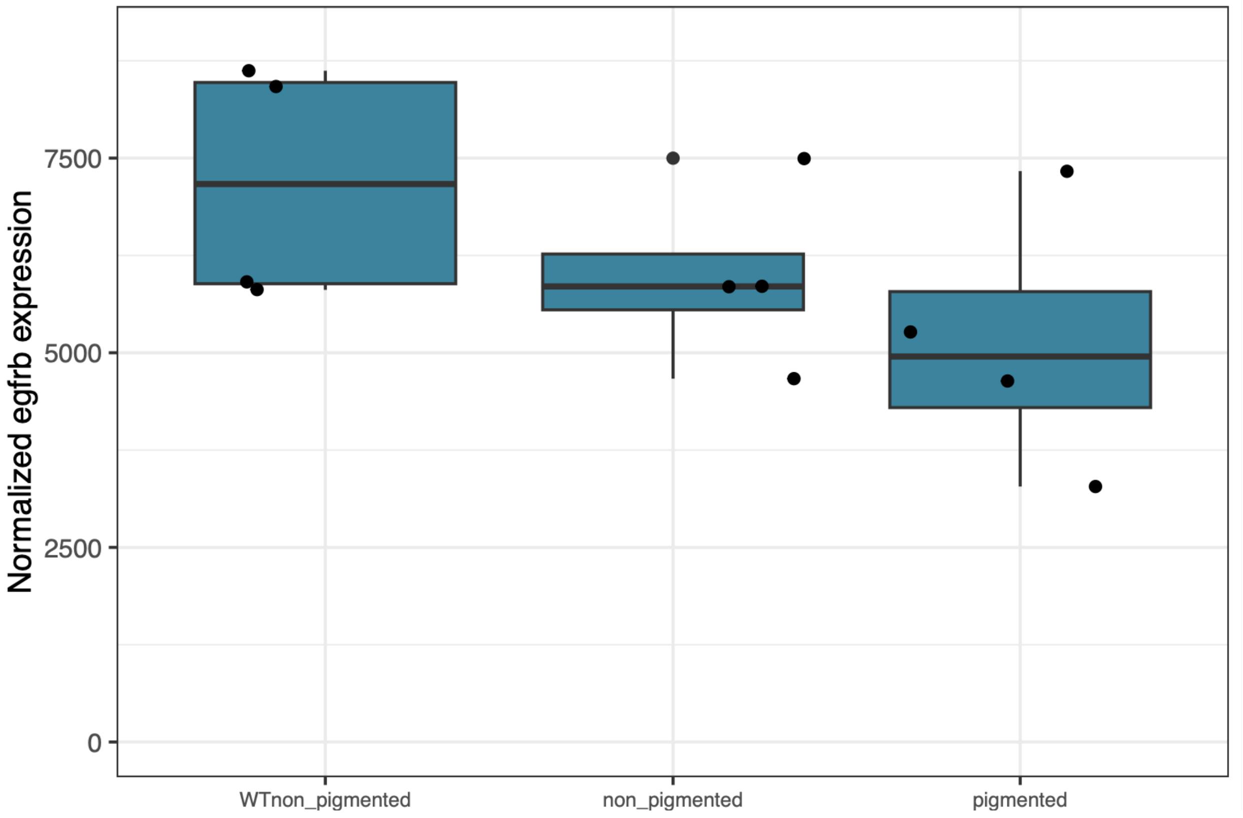


**Figure S12: Normalized expression of *egfrb* across tissues and phenotypes (A)** Normalized expression of *egrfb*, the paralog of *xmrk*, across the 3 different tissue types: wildtype caudal fin tissue, unpigmented tissue from confetti individuals, and pigmented caudal fin tissue. *Egfrb* was not differentially expressed between pigmented confetti tissue and unpigmented tissue on a confetti fish ($p = 0.626$) or between pigmented confetti tissue and unpigmented tissue from an unspotted wildtype fish ($p = 0.798$).

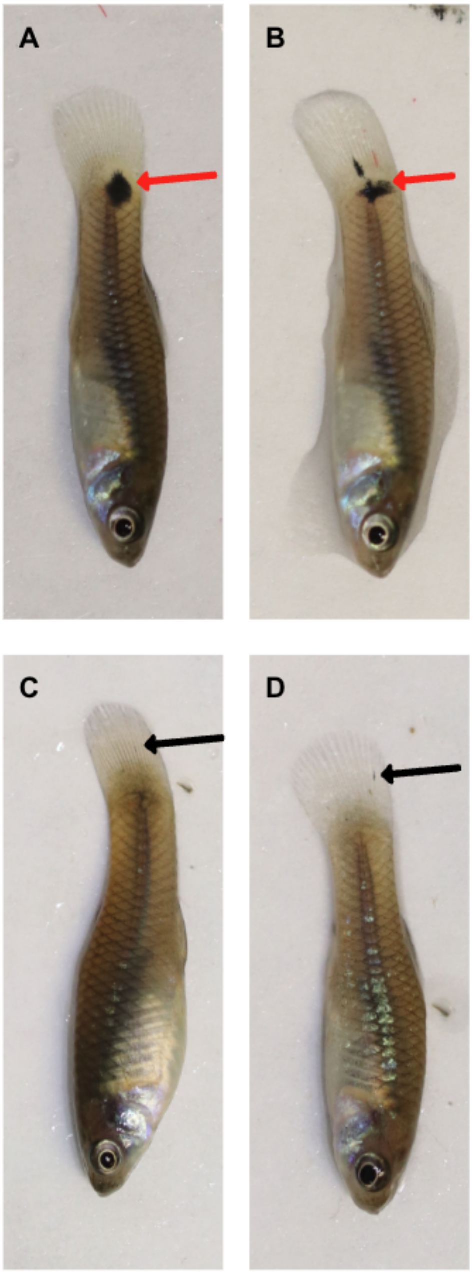


**Figure S13: Representative phenotypes of *X. malinche* × *X evelynae* F1 hybrids relative to pure *X. evelynae*.**
**(A,B)** *X. evelynae* × *X. malinche* F1 hybrid juveniles (age 58 days) with macromelanophore spots on the caudal peduncle (red arrows). **(C,D)** *X. evelynae* juveniles (age 59 days) with small, developing confetti caudal spots and macromelanophores on their caudal fin (black arrows).

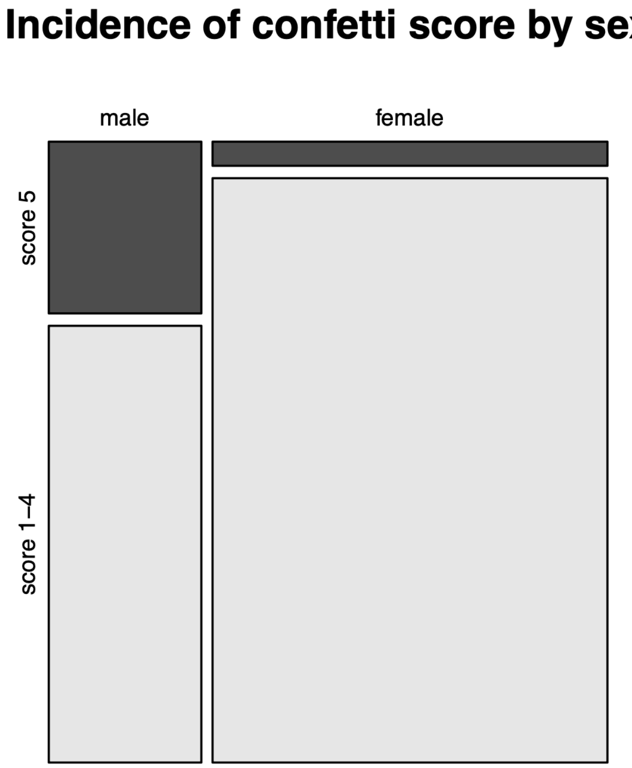


**Figure S14: Males are overrepresented among individuals with the most extreme confetti phenotypes**
Mosaic plot showing proportion of adult males and adult females (juveniles excluded) with the most extreme confetti phenotype (score 5) vs. less extreme pigmentation (scores 1-4, unspotted fish excluded). The width of the bars indicates the number of males and females. Males were more likely to have an extreme confetti phenotype (Fisher’s exact test: $p = 0.000138$).

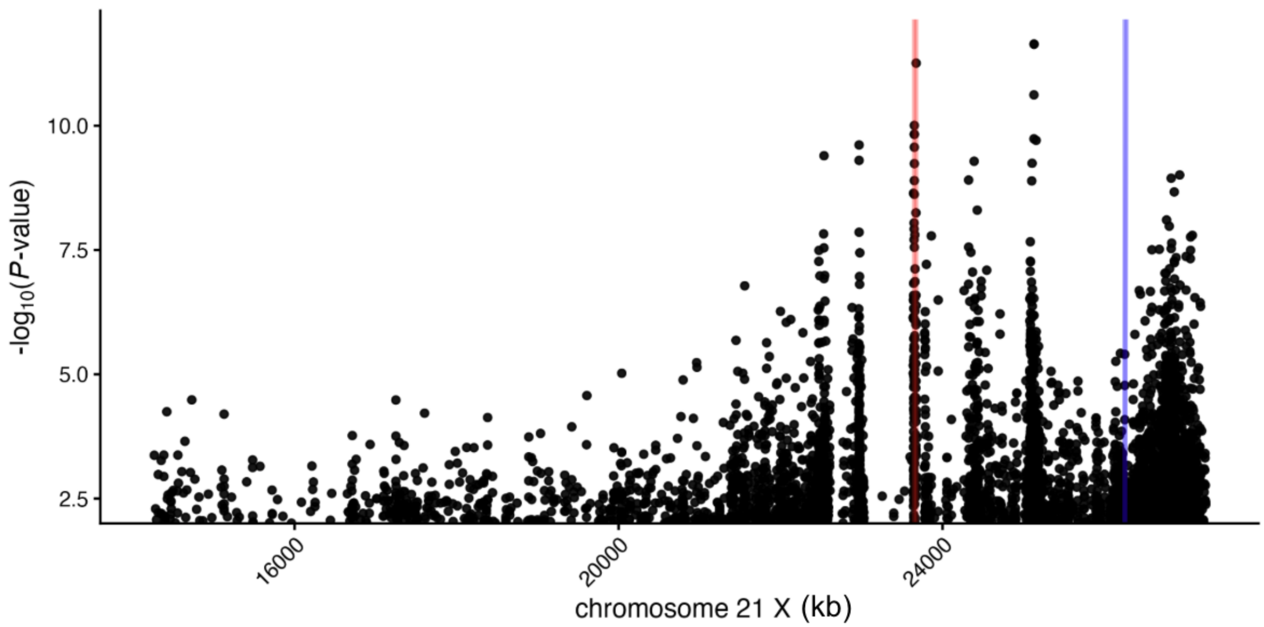


**Figure S15: GWAS results without applying a minimum coverage filter show a peak that colocalizes with the location of *xmrk***

Results presented in the main text include a minimum coverage filter of 80 reads. Here, we plot the unfiltered GWAS results for the distal end of chromosome 21. The red line marks the location of *xmrk,* the blue line marks the location of *xmrk*'s paralog *egfrb.*

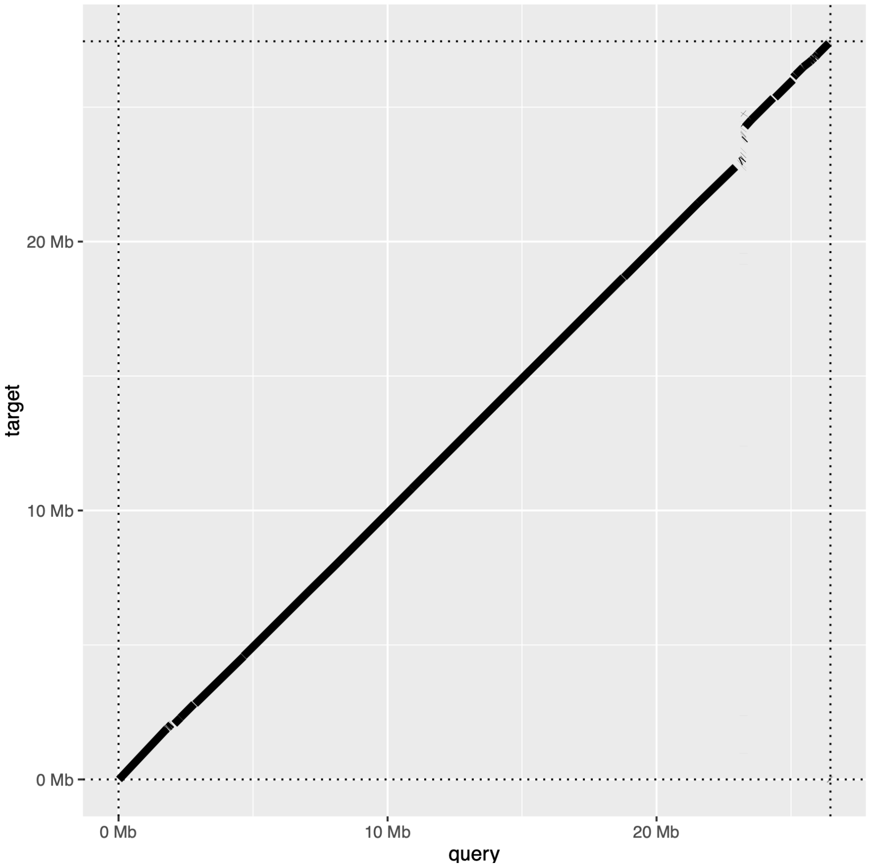


**Figure S16: Alternative visualization of the alignment of chromosome 21-X from an individual with the confetti phenotype and an unspotted, wildtype *X. evelynae***
Chromosome 21 from a long-read assembly of an *X. evelynae* individual with confetti caudal (Y axis) aligned to chromosome 21 from a long-read assembly of an unspotted wildtype individual (X axis), visualized with the pafr R package (David Winter *et al.* 2020) shows a large insertion/deletion structural variant.

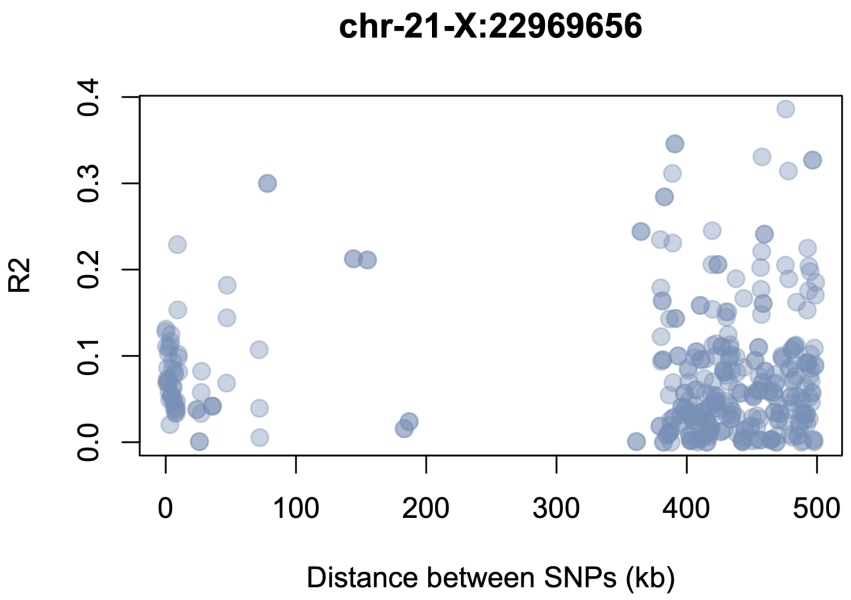


**Figure S17: Linkage disequilibrium in the *xmrk* region.** Average R2 in sliding 500 bp windows as a function of distance to the peak SNP in the GWAS region (near 22.97 Mb) inferred from pseudohaploid data processed in PLINK. Note elevated linkage disequilibrium, sparse data, and lack of monotonic decay in linkage disequilibrium over physical distance in this region of the genome.

**Supplemental tables**

**Table S1: Measurements of macro- and micro-melanophores from images of confetti caudal quantified from photos taken of the caudal fins from four individuals.**

| individual | Cell | Melanophore location | Diameter 1 (µm) | Diameter 2 (µm) | Diameter 3 (µm) | Average diameter (µm) |
|---|---|---|---|---|---|---|
| 1 | 1 | confetti | 124.0 | 50.5 | 99.4 | 91.3 |
| 1 | 2 | confetti | 58.0 | 90.8 | 78.1 | 75.6 |
| 1 | 3 | confetti | 63.3 | 62.5 | 66.7 | 64.2 |
| 1 | 4 | non-confetti | 29.9 | 8.6 | 17.2 | 18.5 |
| 1 | 5 | non-confetti | 23.2 | 24.5 | 11.7 | 19.8 |
| 1 | 6 | non-confetti | 21.4 | 11.5 | 24.3 | 19.0 |
| 2 | 1 | confetti | 356.8 | 147.0 | 244.5 | 249.4 |
| 2 | 2 | confetti | 140.7 | 83.9 | 99.0 | 107.8 |
| 2 | 3 | confetti | 238.3 | 114.2 | 187.1 | 179.8 |
| 2 | 4 | non-confetti | 89.2 | 47.6 | 64.4 | 67.1 |
| 2 | 5 | non-confetti | 47.7 | 33.8 | 42.6 | 41.4 |
| 2 | 6 | non-confetti | 33.9 | 28.7 | 25.0 | 29.2 |
| 3 | 1 | confetti | 77.4 | 49.7 | 57.0 | 61.4 |
| 3 | 2 | confetti | 32.3 | 71.3 | 23.9 | 42.5 |
| 3 | 3 | confetti | 71.7 | 81.8 | 91.3 | 81.6 |
| 3 | 4 | non-confetti | 38.2 | 14.6 | 35.9 | 29.6 |
| 3 | 5 | non-confetti | 44.6 | 21.8 | 34.7 | 33.7 |
| 3 | 6 | non-confetti | 48.5 | 51.9 | 42.3 | 47.5 |
| 4 | 1 | confetti | 102.9 | 77.8 | 81.2 | 87.3 |
| 4 | 2 | confetti | 107.2 | 72.6 | 92.6 | 90.8 |
| 4 | 3 | confetti | 115.1 | 88.5 | 102.4 | 102.0 |

| | | | | | | |
|---|---|---|---|---|---|---|
| 4 | 4 | non-confetti | 53.2 | 50.0 | 30.5 | 44.5 |
| 4 | 5 | non-confetti | 73.2 | 14.9 | 49.5 | 45.9 |
| 4 | 6 | non-confetti | 114.1 | 19.7 | 59.7 | 64.5 |

**Table S2: Genes under the GWAS peak with significant differential expression between pigmented confetti tissue and a wildtype, unspotted fish**

| **Gene ID** | **Gene name** | **Ensembl ID** | **Log fold change** | **Adjusted p-value** |
|---|---|---|---|---|
| g22139 | eif5a | ENSXMAT00000026820.1 | -0.389 | 6.26e-04 |
| g22133 | eif2s3 | ENSXMAT00000011927.2 | -0.453 | 7.39e-04 |
| g22052 | Cdh6 | ENSXMAT00000014211.2 | 0.857 | 1.74e-02 |
| g22056 | mc4r | ENSXMAT00000034758.1 | 3.09 | 3.83e-08 |
| g22063 | xmrk | ENSXMAT00000008846.2 | 5.05 | 1.06e-06 |
| g22064 | Uncharacterized protein | | 3.27 | 6.72e-04 |
| g22150 | B4GALT2 | ENSXMAT00000040096.1 | -1.01 | 1.68e-02 |
| g22170 | Scn5a | ENSXMAT00000011688.2 | 2.78 | 2.20e-03 |

| | | | | |
|---|---|---|---|---|
| g22055 | Drosha | ENSXMAT00000024542.1 | 2.64 | 3.44e-03 |
| g22140 | rpl22l1 | ENSXMAT00000012087.2 | -0.311 | 4.33e-03 |
| g22061 | pol | ENSXMAT00000034964.1 | 5.00 | 7.16e-06 |

**Table S3: Module name, p-value, and correlation with confetti and wildtype phenotype produced through coexpression clustering analysis**

| module | *p*-value: pigmented | correlation: pigmented | *p*-value: WT | correlation: WT |
|---|---|---|---|---|
| MEskyblue2 | 0.287 | -0.335 | 0.031 | 0.622 |
| MEsaddlebrown | 0.165 | -0.428 | 0.018 | 0.667 |
| MEblue | 0.314 | -0.318 | 0.039 | 0.599 |
| MEyellow4 | 0.487 | -0.222 | 0.045 | 0.586 |
| MEmagenta | 0.018 | -0.665 | 0.015 | 0.678 |
| MEred | 0.172 | -0.422 | 0.030 | 0.624 |
| MEskyblue3 | 0.016 | 0.675 | 0.374 | -0.282 |
| MEyellow | 0.019 | 0.662 | 0.268 | -0.348 |
| MEdarkred | 0.007 | -0.726 | 0.325 | 0.311 |
| MEplum1 | 0.015 | 0.679 | 0.007 | -0.727 |
| MEcoral2 | 0.249 | 0.361 | 0.001 | -0.814 |
| MEnavajowhite2 | 0.216 | 0.385 | 0.004 | -0.767 |
| MEthistle2 | 0.791 | -0.086 | 0.025 | -0.641 |
